# How do we align in good conversation? Investigating the link between interaction quality and multimodal interpersonal coordination

**DOI:** 10.64898/2026.05.09.723997

**Authors:** Marcos E. Domínguez-Arriola, Peter C.H. Lam, Alejandro Pérez, Marc D. Pell

## Abstract

Conversations can feel effortlessly engaging or, conversely, difficult and unrewarding. Multiple factors contribute to the experienced quality and outcomes of a conversation, among them how interlocutors align with each other. The present study investigated speech-to-speech, brain-to-speech, and brain-to-brain coordination as markers of interpersonal alignment, examining their relationship with jointly perceived interaction quality and mutual affinity between conversational partners. Pairs of previously unacquainted participants (dyads) engaged in multiple short, free-form conversations on topics of varying interest while their vocal and neural activity were simultaneously recorded in a dual-EEG (“hyperscanning”) setup. We analyzed interlocutors’ prosodic adaptation, neural speech tracking, and neural coordination during each conversation. At the speech-to-speech level, our findings reveal that partners with more positive mutual impressions became more similar in their volume and voice quality over the course of the experiment session, reflecting greater prosodic convergence. At the brain-to-speech level, we found no reliable effect of interaction quality on neural tracking of unfolding speech within any individual region, although topographical differences suggested relative modulation across scalp sites. Finally, at the brain-to-brain level, our findings show that higher perceived interaction quality enhanced inter-brain relationships across frequency bands (alpha and theta) and temporal dependencies (*concurrent*/near-instantaneous and *recurrent*/listener-lagging), with the strongest effects observed for concurrent alpha-band coupling. These findings suggest that distinct coordination processes are involved in how interlocutors experience an interaction and how they establish relational affinity, casting new light into the mechanisms that make a conversation worthwhile.

## 1. INTRODUCTION

What makes a good conversation? People engage in countless social interactions, some of which are perceived as worth one’s time, while others are not. The perceived quality of a conversation is shaped by multiple factors, including interlocutors’ behavioral and vocal cues, interpersonal liking, the presence of novelty or complexity, and the degree to which conversational and social goals are fulfilled (Domínguez-Arriola & Pell, in press; Guydish & Fox Tree, 2021, 2025; Ravreby et al., 2022). Good conversations are also marked by how interlocutors exchange these signals. In real-time conversation, one person’s overt signals of enjoyment can be rewarding to the other, eliciting corresponding signals that, in turn, are rewarding to the first person, creating a positive feedback loop that shapes conversational experience (Hari & Kujala, 2009). Indeed, when interlocutors are mutually responsive and cooperative, conversations tend to be experienced as smooth, effortless, and engaging, giving rise to the subjective experience of *flow* (Koudenburg et al., 2017).

Thus, coordination with another person has been proposed to function as a relational signal of a positive interpersonal stance: by matching an interlocutor’s behavior (e.g., expressive style), affective cues, or expressed ideas, individuals may (often implicitly) convey liking and engagement in the ongoing interaction (Giles et al., 1991; Lakin & Chartrand, 2003). Humans appear to be sensitive to such alignment cues and to experience them as rewarding (Atzil et al., 2014; Cacioppo et al., 2014; Dumas & Fairhurst, 2021; Gvirts & Perlmutter, 2020; Shamay-Tsoory et al., 2019), thereby fostering positive interpersonal outcomes (Cacioppo et al., 2014; Cohen et al., 2024; Kokal et al., 2011; Templeton et al., 2022). Importantly, this alignment extends beyond behavior and subjective experience to the body and the brain: social partners engaged in coordinated interaction tend to exhibit similar—or complementary—physiological and neural activity (Dumas et al., 2010; Stephens et al., 2010). Accordingly, high-quality social interactions are likely to be marked by a high degree of mutual adaptation and multimodal interpersonal coordination: from overt behavior to the underlying neural activity. This study tests this idea by examining speech-to-speech, brain-to-speech, and brain-to-brain coordination across levels of interaction quality.

### 1.1. Multimodal Coordination in Social Interaction

When people interact, their individual actions are difficult to understand in isolation. In conversation, for example, interlocutors’ behaviors (e.g., what they say and how they express themselves) constitute *joint actions*. These actions are mutually dependent and emerge from an evolving, shared conversational context (Clark, 1996; Garrod & Pickering, 2009). Such reciprocity arises from core processes of common attention, mutual prediction, and co-adaptation, whereby interlocutors continuously anticipate each other’s behaviors and mental states in real time and adjust their own accordingly (Hasson & Frith, 2016; Shamay-Tsoory et al., 2019). On this view, interpersonal coordination emerges as conversational partners align at multiple levels, ranging from low-level expressive features of speech to higher-level representations of the unfolding interaction (Dumas & Fairhurst, 2021; Garrod & Pickering, 2009; Pickering & Garrod, 2004; Stolk et al., 2016). This alignment across multiple levels has been suggested to reduce processing demands and energetic costs, making it an intrinsically desirable—and therefore reward-motivated—state for interaction partners (Gvirts & Perlmutter, 2020; Hoehl et al., 2021; Koban et al., 2019). As a result, alignment between interlocutors is thought to serve a core functional role in social interaction, promoting shared understanding, efficient coordination, rewarding interpersonal outcomes, and, ultimately, prosocial behavior (Giles et al., 1991; Guydish & Fox Tree, 2021; Hove & Risen, 2009; Hu et al., 2022; Kokal et al., 2011; Mogan et al., 2017).

Crucially, these processes are implemented in real time by neural systems that themselves become coupled across interacting individuals (Chidichimo et al., 2025; De Felice et al., 2025; Hasson et al., 2012; Hasson & Frith, 2016; Zada et al., 2025). Research using simultaneous dual-brain recordings (“hyperscanning”) has documented such neural coordination—or *inter-brain relationship* (Pérez & Davis, 2023)—as early as 4 months of age (Hoehl et al., 2025; Nguyen et al., 2023) and even across different species (Kingsbury et al., 2019; Y. Yang et al., 2021; Zhang & Yartsev, 2019). It has been linked to social facilitation processes (Adel et al., 2025; Dikker et al., 2022; Hoehl et al., 2021; Shen et al., 2025; Szymanski et al., 2017), while also being systematically shaped by interactional factors such as partners’ relationship characteristics and the structure and reciprocity of the exchange (Ahn et al., 2018; De Felice et al., 2025; Djalovski et al., 2021; Nguyen et al., 2020; Pan et al., 2018; Schwartz et al., 2024; Speer et al., 2024).

Neural coordination driven by the transmission and processing of the speech signal is expected to exhibit temporal asymmetries, with activity in the listener reliably following that of the speaker (Pérez et al., 2022). Accordingly, the temporal profile of neural alignment constrains its functional interpretation: *recurrent* (listener-lagging) inter-brain relationships are consistent with sequential processing of the speech signal, whereas *concurrent* (near-instantaneous) inter-brain relationships are more plausibly linked to common attentional states, predictive processing, and the construction of shared representations or “situation models” (Burns et al., 2025; Kelsen et al., 2022; Menenti et al., 2012; Pérez et al., 2022).

Given that concurrent inter-brain relationships are thought to reflect shared attention and opportunities for mutual prediction and adaptation, more reciprocal and dynamically structured interaction contexts are expected to afford greater opportunities for cross-brain dependence (De Felice et al., 2025; Pan et al., 2018). However, very few studies have investigated neural alignment during self-managed, spontaneous conversation that allows natural turn-taking, interruptions, and backchanneling (e.g., “uh-huh”), with a few exceptions, notably in the developmental literature using fNIRS hyperscanning (Hoehl et al., 2025; Nguyen et al., 2020). For example, Nguyen et al. (2020) found that in mother-child pairs interacting during a cooperative tangram puzzle-solving task, neural alignment in bilateral prefrontal and temporoparietal areas was associated with higher behavioral reciprocity (e.g., smooth turn-taking, as rated by trained coders) and resulted in faster puzzle completion. The authors interpreted neural coordination as a biomarker of interaction quality; interactions that were well coordinated and resulted in enhanced cooperative outcomes (and were therefore high in quality) were characterized by a greater inter-brain relationship. This work suggests that neural coordination indexes the degree to which interaction partners successfully align toward shared goals. Naturalistic conversation constitutes a fundamentally cooperative activity (Bara, 2010), requiring continuous mutual prediction, adaptation, and turn coordination. Yet, how variation in conversational quality maps onto inter-brain relationships remains largely unexplored.

Variation in interpersonal alignment is also reflected in explicit communicative behavior. In conversation, one prominent form of behavioral coordination is *prosodic adaptation*, whereby speakers adapt aspects of their vocal expression to their partner’s (Kruyt et al., 2023; Pardo, 2013). Prosodic adaptation has been suggested to be an intrinsic component of natural dialogue (Giles et al., 1991). For instance, Levitan & Hirschberg (2011) examined a corpus of recorded conversations in which dyads, who could not see each other, played a series of cooperative computer games requiring verbal communication, and quantified prosodic adaptation across four acoustic dimensions that are highly perceptible and play a key role in social voice perception: intensity, fundamental frequency (F0), vocal quality (harmonics-to-noise ratio, shimmer, and jitter), and speaking rate. Across intensity and voice-quality parameters, the authors found that partners spoke more similarly to each other in the second than in the first half of each session’s first game, indicating *convergence* at the level of one communication event (i.e., over the course of a ∼2-minute-long cooperative game). They also observed a significant reduction in F0 distance between the second and first half of the whole experiment, suggesting F0 convergence at the session level (i.e., over the course of ∼45-minute-long recordings). Finally, participants exhibited significant turn-by-turn *synchrony* (shared co-fluctuations) across all acoustic parameters, but particularly voice intensity. Comparable patterns have emerged across a range of speaking contexts (e.g., Levitan et al., 2015; Xia et al., 2014), using diverse analytic methods (e.g., Bonin et al., 2013; Pardo et al., 2013), and targeting different acoustic features (e.g., Edlund et al., 2009; Ostrand & Chodroff, 2021). These findings indicate that interlocutors coordinate their speech production and come to speak more similarly over time.

Prosodic adaptations, much like neural coordination, are systematically shaped by contextual properties of the interaction, including the language spoken, speakers’ roles, cognitive load, personality traits, and the speakers’ relational intent (Abel & Babel, 2017; Giles et al., 1991; Levitan et al., 2015; Michalsky et al., 2018; Natale, 1975; Reichel et al., 2018; Weise et al., 2019). For example, Natale (1975) found that dyads with jointly higher social desirability scores, indexing a shared need for social approval, exhibited greater convergence in mean vocal intensity across three weekly one-hour unstructured conversations. In another study, Michalsky et al. (2018) analyzed 100 dating conversations between previously unacquainted heterosexual participants paired in opposite-sex dyads. Participants rated their overall impression of each conversation on a 10-point Likert scale. The results showed that, for female speakers, impressions of the conversation’s quality significantly predicted the extent to which their F0 converged toward that of their partner. Overall, the existing evidence points to a complex interplay between individual differences and situational demands in shaping prosodic adaptation.

Taken together, this body of work points to interpersonal alignment as a multimodal and context-sensitive process through which interaction partners coordinate to produce joint social action, linking subjective conversational experience to both prosodic and inter-brain relationships (Dippong & Overton, 2025). This is in accordance with the interactive-alignment model, which predicts alignment across multiple representational levels during dialogue (Garrod & Pickering, 2009; Pickering & Garrod, 2004), expressed as coordination across behavioral and physiological channels (Menenti et al., 2012), as well as with Communication Accommodation Theory (Giles et al., 1991, 2023), which highlights the context dependency and relational consequences of interpersonal (dis)alignment. However, it remains unclear how behavioral (e.g., prosodic) and neural coordination jointly unfold during naturalistic, truly social interaction.

### 1.2. Neural Tracking of Speech Rhythms

In addition to speech-to-speech and brain-to-brain coordination, alignment can also be observed at the brain-to-speech level through the neural tracking of speech rhythms. Perceptual rhythms are pervasive in spoken communication (e.g., at the syllabic and suprasegmental levels) and are leveraged by the brain to parse and process connected speech (Ding et al., 2016; Poeppel & Assaneo, 2020). In particular, slow neural oscillations in a listener’s brain “entrain” to these pseudorhythmic amplitude modulations in speech, a process known as neural speech tracking (NST), which can be measured with EEG and provides a quantitative index of speech processing (Brodbeck & Simon, 2020; Luo & Poeppel, 2007; Pérez et al., 2022). Neural tracking of continuous speech is influenced both by bottom-up acoustic and top-down motivational features of the perceived stimulus (Decruy et al., 2020; Fiedler et al., 2019; Holtze et al., 2021; Lesenfants & Francart, 2020; Obleser & Kayser, 2019; Pascucci, 2025; Vanthornhout et al., 2019). A recent study showed that NST in frontal channels was contextually modulated by the speaker’s prosody in conversation-like stimuli, such that an “engaging” speaking style enhanced listeners’ neural tracking of social anecdotes, but only when their content was interesting (Domínguez-Arriola et al., under review). During naturalistic dialogue, interlocutors attend to and track each other’s communicative rhythms (Pérez et al., 2017, 2019). Because NST provides a quantitative measure of online speech processing with a characteristic stimulus-response lag (∼110 ms), it has been proposed as an indirect proxy of speech-driven contributions to recurrent inter-brain relationships (see Pérez et al., 2022). Thus, examining real-time NST alongside other forms of interpersonal coordination will provide insight into the possible contribution of speech processing to interpersonal alignment during naturalistic dialogue.

### 1.3. The Present Study

Leveraging EEG hyperscanning during naturalistic conversation, this study investigated whether the jointly perceived quality of an interaction enhances interpersonal coordination at three levels: speech-to-speech (prosodic adaptation), brain-to-speech (NST), and brain-to-brain (inter-brain relationship). To this end, we simultaneously recorded EEG and audio data from pairs of participants who had never met before as they engaged in a series of short, spontaneous conversations in which they were co-present but visually occluded from each other. Visual occlusion helps preserve the conversation’s naturalistic quality while constraining the interactional signal to the vocal domain (Pérez et al., 2017, 2019). Partners also did not see each other until the end of the experiment, preventing visual cues and physical appearance from influencing social evaluations. To experimentally modulate the inherent value of individual conversations, dyads discussed topics that were jointly rated as either highly interesting or uninteresting (Domínguez-Arriola & Pell, in press). The conversations were fully free-form, allowing natural interruptions, overlaps, and backchannels, and each speaking turn was subsequently delineated from the audio recordings. This design allowed participants to engage not only with the conversational content but also in natural turn-taking dynamics, which are a central component of real-world social interaction (Kawasaki et al., 2013; Templeton et al., 2022). Participants confidentially provided judgements about each four-minute conversation and, at the end of the session, about their mutual social impressions.

Prosodic adaptation was examined for intensity, fundamental frequency (F0), harmonics-to-noise ratio (HNR), and speaking velocity (Levitan & Hirschberg, 2011). Similarly to Levitan & Hirschberg (2011), we quantified acoustic-prosodic *convergence* at both the level of the whole experiment session and, more locally, at the single-conversation level, and *synchrony* within conversations. We expected dyads to adapt prosodically at the conversation level, manifesting both convergence and synchrony, with higher interaction quality enhancing both forms of mutual adaptation (Michalsky et al., 2018). We also hypothesized that prosodic convergence across the entire testing session would be predicted by interlocutors’ session-level social impressions of one another.

Across the NST and neural coordination literatures, statistical dependence between signals has been quantified using a wide range of approaches (Gross et al., 2021; Hakim et al., 2023). More recently, there has been a growing call to adopt information-theoretic metrics, as these are sensitive to non-linear dependencies and can capture relationships that are complementary rather than strictly mirrored (Burns et al., 2025; Chidichimo et al., 2025; Hasson & Frith, 2016). One candidate approach is the Gaussian Copula Mutual Information framework (GCMI). This method isolates the empirical copula between two variables via rank transformation, transforms the marginals to standard normal distributions, and then estimates mutual information using a Gaussian parametric formulation. The resulting GCMI values are robust to outliers and provide a lower-bound estimate of the true mutual information (see Ince et al., 2017). Here, we employ this metric to compute the statistical dependence between EEG and speech envelope signals (i.e., NST; e.g., Pérez et al., 2022, 2026) and between EEG signals across interacting brains (i.e., inter-brain relationships). At the brain-to-speech level, we hypothesized that interaction quality would positively modulate GCMI-based NST estimates—hereafter, GCMI_NST_—particularly in frontal regions (Domínguez-Arriola et al., under review).

EEG-based examination of inter-brain relationships has focused on different frequency ranges, which are often associated with different functional roles (Angioletti et al., 2024; Djalovski et al., 2021; Dumas et al., 2010; Pérez et al., 2017). In particular, alpha-band activity (8-12 Hz) has been proposed to track fluctuations in joint attention during social interaction (Dikker et al., 2017; Pérez et al., 2017, 2019). By contrast, neural coordination in the theta band (3-7 Hz) may be linked to similar linguistic and paralinguistic processing (Pérez et al., 2017) and to shared behavioral rhythms that support interpersonal coordination during cooperation and joint action (Wang et al., 2020; M. Yang et al., 2023). Here, we index GCMI-based *concurrent* and *recurrent* inter-brain relationships (GCMI_IB_) in the alpha and theta frequency bands. These bands capture key joint attentional, linguistic, and social processes (Kingsbury & Hong, 2020; Pérez et al., 2017, 2019), and are generally less susceptible to movement- and speech-related artifacts than lower-frequency delta activity or higher-frequency beta rhythms, respectively (Goncharova et al., 2003). We hypothesized that higher interaction quality would be associated with stronger inter-brain relationships across frequency bands, reflecting increased attentional, (extra)linguistic, and representational alignment.

Together, this multimodal hyperscanning approach tests a unified account of interpersonal alignment. By examining how interaction quality shapes prosodic adaptation, neural speech tracking, and inter-brain relationships within the same dyadic exchanges, the present study constitutes a comprehensive investigation of how subjective interaction value modulates the shared processes that support naturalistic social communication.

## 2. METHODS

### 2.1. Participants

Sample size was determined a priori using effect sizes from prior studies using EEG hyperscanning in interactive paradigms (e.g., Bevilacqua et al., 2019; Djalovski et al., 2021; Pérez et al., 2017; Schwartz et al., 2024). We powered the study to detect effects of *d* > 0.7 with 90% power at α = 0.05 (two-tailed), which yielded a target sample of 24 dyads. Fifty participants (25 unacquainted dyads; 26 women, 22 men, 2 non-binary; *M* age = 22.4, SD = 4.3) took part in the study. Data from one dyad were excluded from the neural analyses due to a technical issue, yielding a final sample of *n* = 48 (24 dyads) for neural analyses and *n* = 50 (25 dyads) for behavioral/acoustic analyses. Same-sex pairings were determined based on participants’ reported sex at birth. All participants were native speakers of North American English with no reported history of speech, hearing, psychiatric, or neurological disorders. Informed consent was obtained before participation, and participants received $50 CAD in compensation for 2.5 hours. The study was approved by the Research Ethics Board of the Faculty of Medicine and Health Sciences at McGill University (study number A05-B22-01A).

### 2.2. Topic Interest Screening

Given that the semantic content of a conversation is a key determinant of its perceived value (Domínguez-Arriola & Pell, in press), we introduced variation in the intrinsic value of different conversations via topic selection. Specifically, upon signing up for the experiment, participants completed a questionnaire in which they rated their interest in 119 different topics (e.g., cooking, golf, the Oscars, astrology) on a 6-point Likert scale ranging from “I really don’t care about this” to “I am very passionate about this”. Dyads were formed by matching participants who shared six topics both rated as highly interesting (scores of 5 or 6) and six topics both rated as highly uninteresting (scores of 1 or 2), without participants’ knowledge of this matching procedure. This process ensured that conversation partners shared similar (dis)interests and similarly valued different conversation topics.

### 2.3. Experimental Setting

Each participant was greeted separately and escorted to the testing booth, where the two members of the dyad sat separated by a curtain. Dyad members were unacquainted and did not see each other at any point until after the experiment was completed, ensuring that social evaluations were based solely on vocal communication. Two dyads, however, saw each other briefly before the experiment because they arrived at the testing site simultaneously. The experiment was conducted in a large, sound-attenuating booth, with participants seated side by side (separated by the curtain) facing a computer monitor (see Figure 1). A microphone (Yeti, Logitech for Creators; 16-bit/48 kHz) was positioned in front of both participants to record their vocal productions. Presentation of all instructions and stimuli was controlled via a custom program developed with the Psychophysics Toolbox, Version 3 (Brainard, 1997; Kleiner et al., 2007). EEG amplifiers and other equipment were placed on a table behind the participants. The study was self-paced, with participants initiating each trial using a keyboard. Experimenters monitored the session from outside the booth, observing through a rear window, listening through headphones, and tracking EEG signals on external monitors.

**Figure 1:**
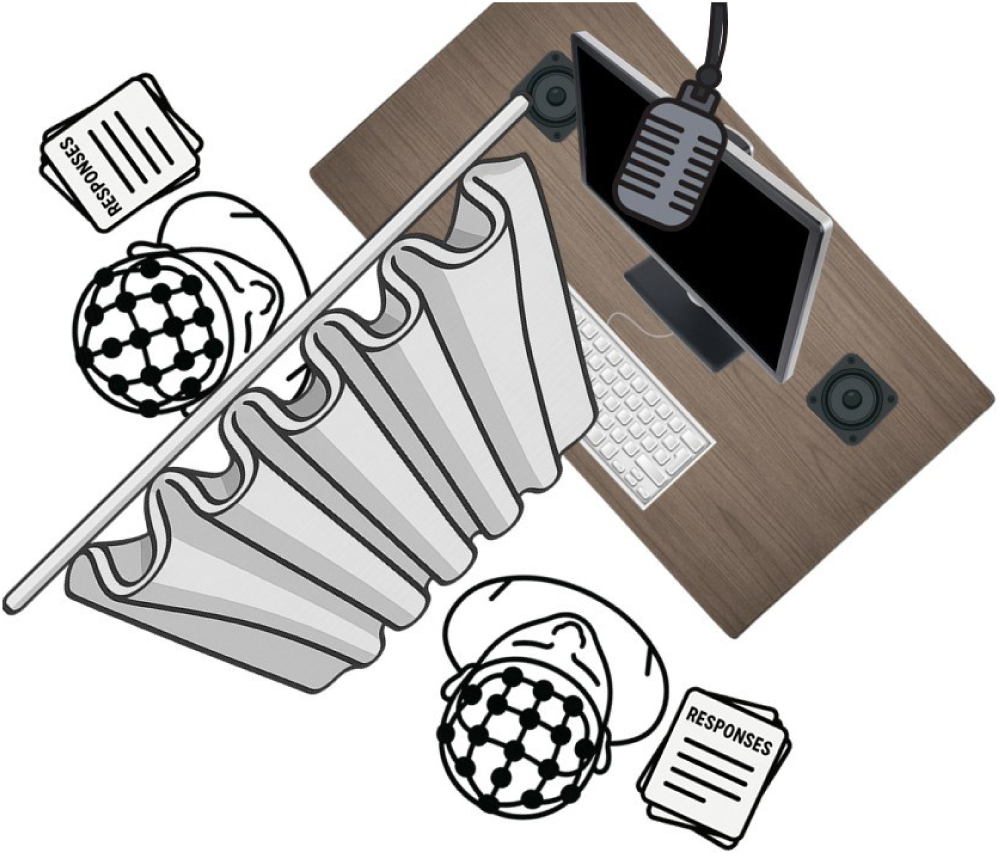
Hyperscanning setup. Conversation partners were separated by a curtain and never saw each other until the end of the experiment.

During EEG setup, participants were instructed not to talk to each other until the start of the experiment, though they could communicate with the experimenters for questions or requests. They were told that they would take part in a series of conversations with one another and were provided with a list of discussion topics and optional prompts, allowing them to prepare in advance if desired. This step was included to minimize prolonged silence, particularly for topics that participants found uninteresting. After EEG setup, participants received detailed instructions. They were asked to engage in natural conversations about the topics shown on screen and to sustain the dialogue for the full duration of each trial. Participants were further instructed to remain relaxed and still, speaking naturally while looking at the fixation cross on screen, minimizing gestures, large body movements, and intense laughter.

### 2.4. Task and Procedure

The experimental task comprised 12 four-minute conversations on topics preselected to differ in their perceived value to participants (see Topic Interest Screening). Each trial began with a slide providing brief instructions (e.g., “In this trial you will talk about…”). The task started after both participants pressed a button to indicate readiness. A screen indicator showed whether each participant had already pressed the button, allowing brief self-paced breaks between trials without verbal communication. Following confirmation from both participants, the screen presented a short countdown (3, 2, 1), after which a fixation cross appeared and remained onscreen for the full trial duration. During conversation trials, the fixation cross changed color 20 seconds before the end of the trial to signal participants to gradually conclude their discussion. Participants were instructed to speak as naturally as possible while maintaining their gaze on the fixation cross and to try to contribute equally to the conversation until the end of the trial.

After each conversation, participants confidentially answered four questions in a response booklet placed beside them: i) *How much longer would you be willing to continue this conversation? (any time unit)* This open-ended item is an adaptation of the time-bidding task, indexing perceived social reward value through participants’ willingness to invest additional time in a social stimulus or situation (Domínguez-Arriola & Pell, in press). ii) *How interested were you in this topic?* This question sought to assess their perceived interest in the topic at the time of the conversation on a 6-point Likert scale. iii) *How effortful was this interaction?* It has been suggested that successful social interactions feel less effortful (Koudenburg et al., 2017; Wheatley et al., 2012), making this a suitable indicator of interaction quality. This item used a 7-point Likert scale. iv) *What was the topic you just discussed?* This open-ended item served as a simple check to ensure that participants remained on topic and completed the correct page of the response booklet.

The session was structured into six blocks, each including one high-interest and one low-interest conversation in random order. Each block also included a 2-minute silence trial, during which participants fixated on a cross without speaking, and a 2-minute listening trial, in which they heard a pre-recorded conversation from same-gender pilot dyads. These conditions are not analyzed here, as they fall outside the scope of the present research questions. The first block additionally included one practice conversation for participants to become comfortable with the format (Figure 2). After completing the six experimental blocks, participants confidentially completed a questionnaire adapted from a previously employed set of social implication scales (Montepare et al., 2014; see Appendix A). The questionnaire assessed impressions of the partner’s approachability, friendliness, competence, and attractiveness, along with participants’ willingness to develop a closer personal or professional connection. Before completing the questionnaire, participants were reminded that their responses were fully anonymized and would not be accessible to their partner or identifiable by the research team.

**Figure 2:**
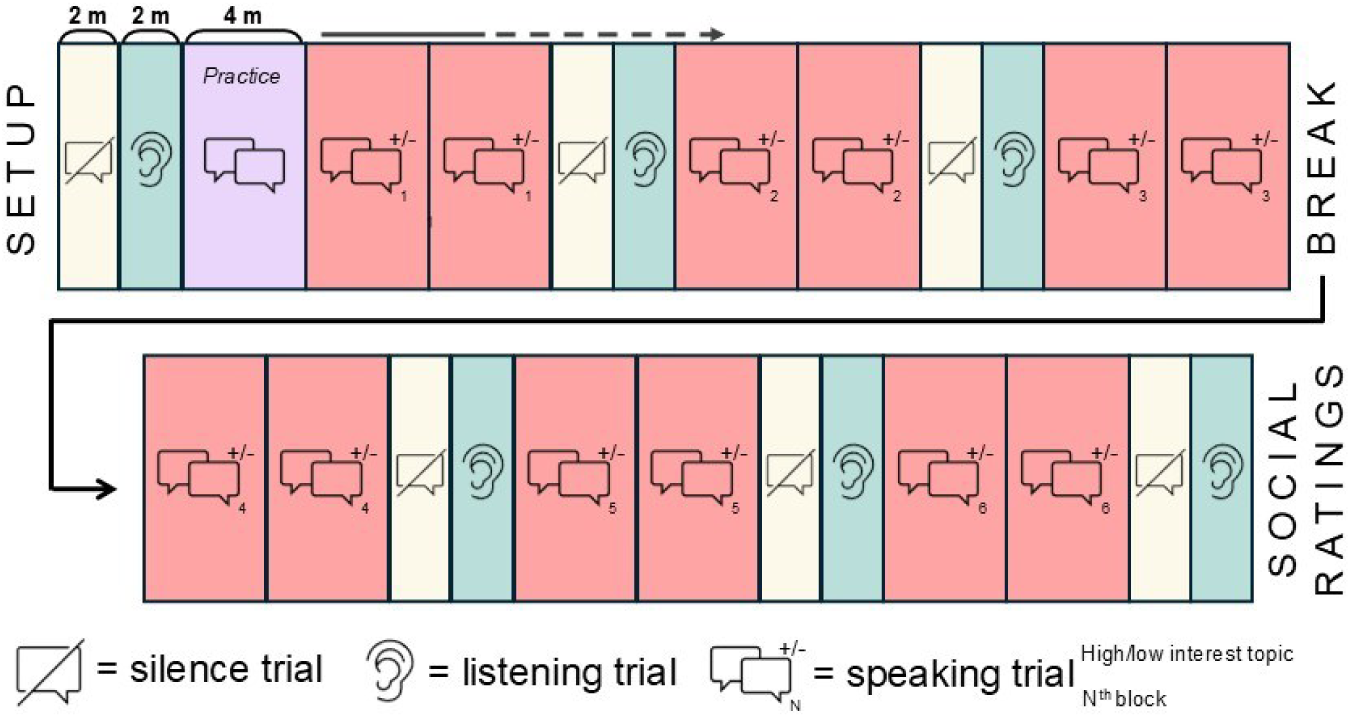
Trial sequence. The session consisted of six blocks, each including two speaking trials, one silence trial, and one listening trial. Block 1 additionally included a practice trial. After completing all blocks, participants provided confidential social ratings of their partner.

### 2.5. EEG Hyperscanning Recording

EEG signals from each subject were recorded using a 32-channel Ag/AgCl actiCAP snap system and LiveAmp amplifier (Brain Products, Germany) at a sampling rate of 500 Hz.

Electrodes were positioned according to the international 10-20 system with individual Reference (FCz) and Ground (AFz) electrodes. Synchronization between amplifiers was achieved through hardware trigger forwarding, allowing offline alignment of recordings with near-millisecond precision (Kreilinger, 2022). The microphone’s signal was synchronously recorded along with the EEG via a StimTrak device (Brain Products, Germany). Impedance measurements were kept below 10 kΩ and signals were recorded and continuously monitored using BrainVision Recorder. EEG markers were time-locked to the beginning and end of each trial.

### 2.6. Data Preprocessing

#### 2.6.1. Behavioral Data Preprocessing

Participants rated their willingness to extend the interaction (time bids), perceived effort, and topic interest after each conversation trial. Time bids were open-ended and therefore expressed on individual-specific scales of time investment as a function of reward value; these responses were therefore normalized for cross-participant comparability. Principal component analysis (PCA) with these three ratings revealed a primary component explaining 71.5% of variance, with strong positive loadings for time bids (0.60) and topic interest (0.60), and a negative loading for perceived effort (-0.53). This component acts as a weighted summary of the three variables and will be hereafter referred to as *perceived interaction quality* (PIQ), reflecting dyads’ engagement with and overall impression of each interaction.

Participants also rated their conversational partner on 12 scales assessing perceived social attributes at the end of each experimental session (Appendix A). PCA revealed a primary component explaining 42.4% of variance, with substantial loadings from “would make me feel at ease” (0.36), approachability (0.32), friendliness (0.31), “would like to work with them” (0.30), attractiveness (0.29), and competence (0.26), among others. Notably, this component encompassed both warmth- and competence-related constructs, suggesting it reflects a general social approach-or-avoid tendency (Cuddy et al., 2008). We refer to each dyad’s mean score on this component as *mutual affinity*.

#### 2.6.2. Audio Preprocessing

Continuous audio from each experimental session was manually divided into 12 four-minute trials. Google Cloud Speech-to-Text (model: *chirp_2*; language: *en_US*) was then used to extract word-level onset and offset times relative to each trial’s onset. We then inspected the output, manually corrected transcription errors, and diarized each segment. Because timing information was unreliable during overlapping speech, we also manually adjusted the onset and offset of each speaking turn using the audio waveform and spectrogram. A speaking turn was defined as a continuous period during which one participant holds the floor, beginning at the onset of that participant’s speech and ending when their speech stops, either before the interlocutor takes the floor or after the interlocutor has already begun speaking, thus allowing consecutive turns to overlap (Sacks et al., 1974). Finally, the amplitude envelope of each speech excerpt was extracted using the mTRF MATLAB toolbox (Crosse et al., 2016), which computes the signal’s root-mean-square energy over time and applies a logarithmic intensity scaling (Lalor & Foxe, 2010).

The GeMAPS acoustic feature set (Geneva Minimalistic Acoustic Parameter Set; Eyben et al., 2016) was extracted for each speaking turn of each conversation partner using the openSMILE toolkit. Here, we focused on four representative acoustic parameters that have exhibited prosodic adaptation in previous literature (Levitan & Hirschberg, 2011): fundamental frequency (F0), intensity, harmonics-to-noise ratio, and voicing rate (i.e., voiced segments per second). We chose voicing rate instead of speaking rate as an index of speaking velocity to accommodate non-linguistic utterances (e.g., fillers, laughter) within each speaking turn, which cannot be reliably quantified by syllable count (Jiao et al., 2015; Samudravijaya et al., 1998).

#### 2.6.3. EEG Preprocessing

Figure 3, right panel, illustrates the EEG preprocessing workflow. EEG data were processed using the EEGLAB toolbox (Delorme & Makeig, 2004) through custom scripts. Raw EEG data were initially band-pass filtered between1 and 40 Hz using a fourth-order Butterworth filter to attenuate slow drifts and high-frequency noise. The data from the two partners in each dyad were then merged into a single continuous dataset based on common event latencies (from the forwarded triggers). Next, the Praat-annotated audio recordings were down-sampled and precisely aligned to the EEG data using the point of maximal cross-correlation between the audio and StimTrak signals. This alignment allowed us to add offline event markers to the EEG data at the exact start and end of each speaking turn, based on careful manual annotation, allowing us to examine neural activity separately during periods when each partner was speaking within a conversation.

**Figure 3:**
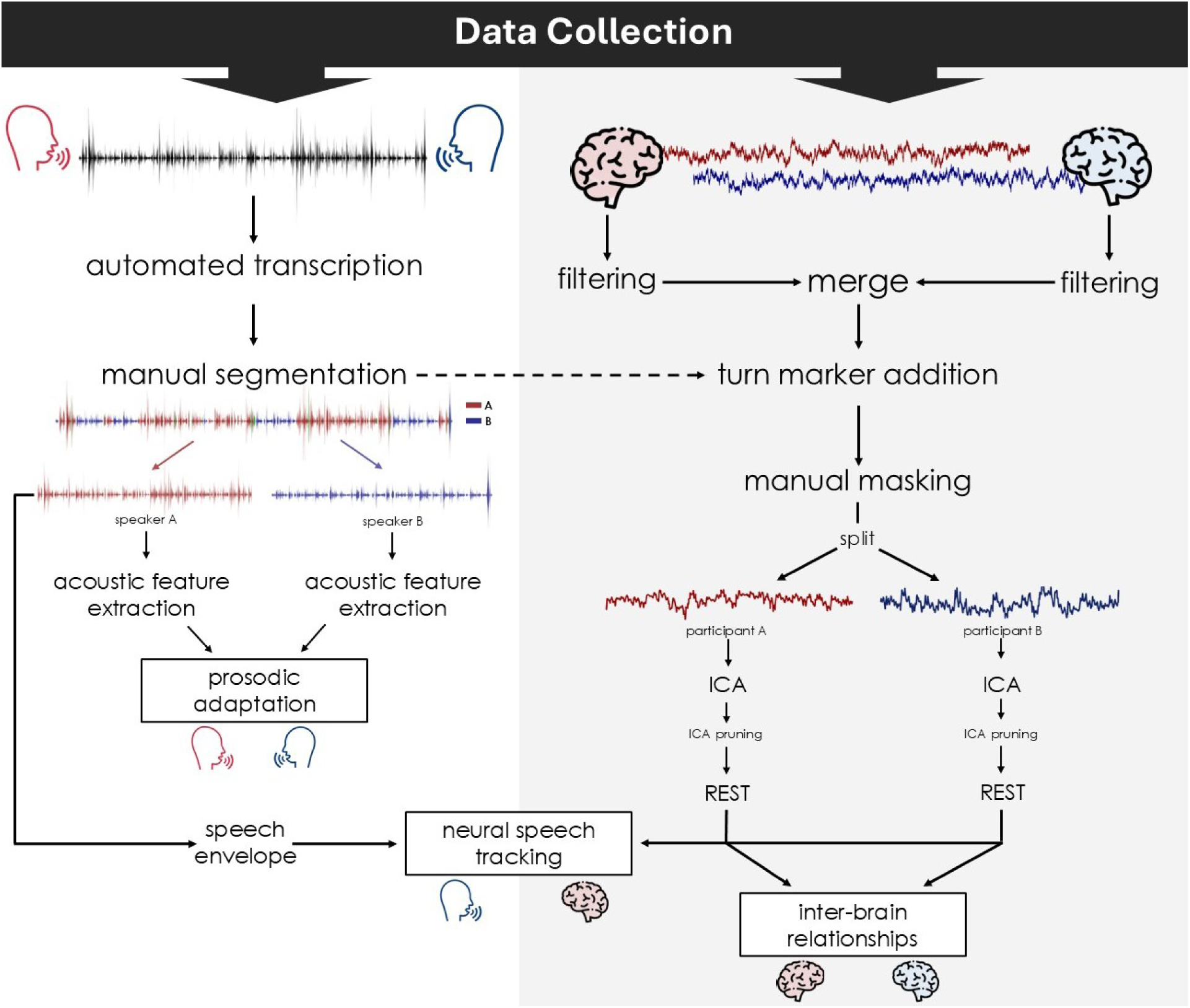
Preprocessing pipeline. Left panel: audio preprocessing steps. Right panel: EEG preprocessing steps.

The EEG data were then visually inspected, and segments exhibiting excessive movement or gross artifacts were manually masked. Since both participants’ signals were merged, this process ensured that identical segments were masked in both datasets. Data segments were not removed at this stage; instead, masks were saved and used to exclude these segments during all downstream analyses. The data were then split again and Independent Component Analysis (ICA) was performed individually for each participant’s data (Delorme et al., 2007; Makeig et al., 2004). Individual components accounting for blinks, speaking-related muscle activity, or line noise were removed from the data. The present combination of band-pass filtering and ICA decomposition has been used to remove most signal fluctuations related to body movement and speaking in naturalistic, dual-EEG paradigms (Pérez et al., 2019; Tamburro et al., 2024). Finally, the EEG signals were re-referenced with the Reference Electrode Standardization Technique (REST), which approximates a neutral reference based on the scalp distribution of potentials, thereby reducing reference-related bias in scalp EEG analysis (Dong et al., 2017; Yao, 2001).

In preparation for the neural speech tracking (NST) analysis, EEG segments in each conversation where a single participant was speaking (i.e., excluding backchannels, between-turn silent gaps, and speaking overlaps) were extracted. Segments under one second were excluded to ensure each excerpt contained some amount of connected speech. For each conversation and participant, we concatenated EEG segments corresponding to intervals in which only the conversational partner was speaking. The StimTrak channel data were replaced with the corresponding speech envelope, precisely aligned via cross-correlation and down-sampled to 500 Hz (Pérez et al., 2022). All channels were then band-pass filtered between 1 and 10 Hz—this range is widely used in neural speech-tracking research to encompass delta and theta activity associated with syllabic and suprasegmental speech encoding (Ding & Simon, 2014; Fiedler et al., 2019; Pérez et al., 2022; Zion Golumbic et al., 2013). The resulting dataset was used for the NST analysis.

In preparation for the neural coordination analysis, we extracted two frequency bands for each conversation and participant using zero-phase FIR filtering: theta (2–8 Hz) and alpha (8–13 Hz). For each conversation trial, the filtered data were segmented into speaker-specific periods using temporal masks derived from the speech-annotation procedure. Speaking turns shorter than 4 s were excluded to ensure that only sustained speaking and listening periods contributed to the analysis. The remaining segments were concatenated separately for each speaker and conversation, yielding two aggregated neural time series per dyad corresponding to periods of speaking and listening for each partner. The resulting dataset was used for estimating and analysing neural coordination between interaction partners (see below).

### 2.7. Analyses

All signal processing and GCMI calculations were conducted in MATLAB (R2023a). All statistical analyses were implemented in R (R Core Team, 2024) with the *lme4* (Bates et al., 2015) and *lmerTest* (Kuznetsova et al., 2017) packages to fit linear mixed-effects models (LMMs). Model selection was theoretically guided and refined through empirical model comparisons based on likelihood ratio tests and information criteria (i.e., AIC and BIC). When a model failed to converge, we first applied an automated refitting procedure that iteratively updated starting values; if convergence was still not achieved, we then removed random-effect correlations and iteratively excluded lower-variance random effects. Where the model included significant interactions, estimated marginal means (EMMs; for categorical × categorical interactions) or estimated marginal trends (EMTs; for continuous × categorical interactions) were computed at relevant combinations of the interacting factors using the *emmeans* package (Lenth, 2025).

#### 2.7.1. Prosodic Adaptation Analysis

Prosodic adaptation was assessed using three complementary approaches, each capturing a distinct aspect of how conversation partners align their vocal expression (Edlund et al., 2009; Levitan & Hirschberg, 2011). We first examined whether conversation partners coordinated their turn-by-turn vocal acoustic modulations (i.e., *conversation-level synchrony*); namely, we asked whether participants’ prosody could be meaningfully predicted from their partner’s prosody in the preceding turn. We also tested whether such synchronization was stronger in conversations characterized by greater PIQ. Turn-level acoustic measures were modeled using a mixed-effects linear regression, including all interactions among the partner’s preceding acoustic measure, PIQ, and acoustic feature. The model included random intercepts and random slopes for both the partner’s preceding acoustic measure and conversational involvement, per partner nested within dyad. In Wilkinson notation:

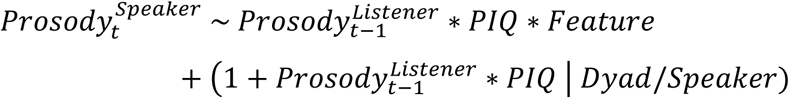

Next, we examined whether conversation partners became more similar over the course of individual conversations (∼ 240 s; i.e., *conversation-level convergence*), and whether this was modulated by Perceived Interaction Quality (PIQ). To do this, we categorized speaking turns within each conversation into temporal stages and focused on Early (0 – 80 s) and Late (160 – 240 s) turns, retaining only segments in which both speakers produced at least two turns at each stage. For each acoustic feature, we computed the distance between conversation partners at the Early and Late stages. This distance was then modeled with mixed-effects linear regression model with all interactions among time, PIQ, and acoustic feature, with random intercepts and random slopes for time and PIQ specified per dyad:

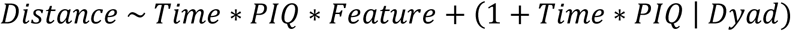

Finally, we assessed whether conversation partners spoke more similarly to each other at the end compared to the beginning of the experiment session (i.e., *session-level convergence*), and whether this convergence was influenced by their mutual affinity. For each acoustic feature (F0, intensity, HNR, and voicing rate) we computed the mean distance between partners in each dyad during the initial (blocks 1 and 2) and final (blocks 5 and 6) segments of the experiment session. We then fit a mixed-effects linear regression model for this distance, as a function of the interaction between time (early vs. late), mutual affinity, and acoustic feature, with random intercepts and random slopes for time and mutual affinity specified per dyad.

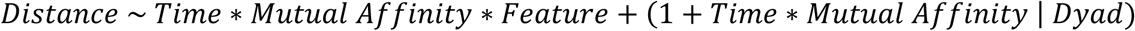

#### 2.7.2. Neural Speech Tracking Analysis

NST was quantified using the Gaussian Copula Mutual Information (GCMI) framework for each conversation and speaker-listener direction using GCMI Toolbox functions (Ince et al., 2017). Specifically, each neural and speech-envelope signal was first copula-normalized. This step transforms each channel’s amplitude distribution to a standard normal marginal while preserving temporal dependencies (Ince et al., 2017). Then, using the normalized signals, the mutual information (i.e., GCMI) between each EEG channel and the corresponding speech envelope was calculated across a range of stimulus-response lags (50 to 300 ms), encompassing canonical latencies for connected-speech neural processing (Fiedler et al., 2019; Pérez et al., 2022). The resulting GCMI_NST_ values were scaled by the duration of each speaker’s signal within each conversation to correct for GCMI’s linear dependence on signal length. For each conversation and speaker-listener direction, NST was defined as the maximum GCMI value within the sampled lag window (50-300 ms).

We then assessed whether the perceived quality of interaction modulates the extent of listeners’ neural speech tracking (i.e., their online neural encoding of connected, real-time speech), while controlling for acoustic covariates that bottom-up influence NST. We used elastic net regression to identify a minimal yet maximally informative set of acoustic covariates to control for in the main statistical model (Tibshirani, 1996; Zou & Hastie, 2005). Starting from a comprehensive set of acoustic parameters potentially relevant to speech encoding (F0 mean and SD, intensity mean and SD, alpha ratio, HNR, turn duration, gap duration, and voicing rate), and listener-derived GCMI_NST_ values as response, an elastic net procedure that tuned α values (0.5-1.0) and constrained the number of retained features yielded four predictors as the most relevant: loudness, loudness variability, F0, and F0 variability.

GCMI estimates are non-negative and typically exhibit a right-skewed distribution due to their zero-bound constraint. In such cases, one may either apply a variance-stabilizing transformation and use standard parametric tests or alternatively rely on generalized linear models tailored to right-skewed, non-negative data (e.g., Gamma GLMs; Hammouri et al., 2020). Inspection of empirical density plots and Q-Q diagnostics in our dataset motivated the use of a square-root transformation for all GCMI response variables prior to statistical modeling.

Transformed GCMINST values were analyzed using an LMM with PIQ and ROI, as well as their interaction, as main predictors. The model further included all selected acoustic covariates as fixed effects. Random intercepts were specified for dyad, listener within dyad, and electrode, and a random slope of PIQ was included for listener within dyad to account for between-listener differences in the association between interaction quality and GCMINST:

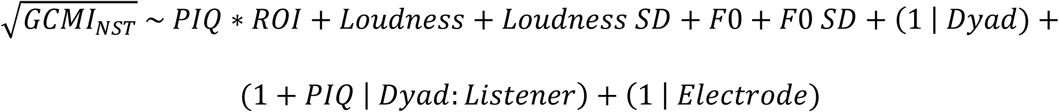

For all neural analyses, we defined six ROIs that provided sufficient spatial specificity to capture effects across frontal, temporal, and central scalp regions: left-anterior (Fp1, F3, F7), medial-anterior (Fz, FC1, FC2, FCz), right-anterior (Fp2, F4, F8), left-temporal (FC5, T7, CP5), medial-central (C3, Cz, C4, CP1, CP2), and right-temporal (FC6, T8, CP6) (Djalovski et al., 2021; Jiang & Pell, 2015; Schwartz et al., 2024).

#### 2.7.3. Neural Coordination Analysis

We estimated cross-brain statistical dependence using the Gaussian Copula Mutual Information (GCMI) framework. First, for each speaker-listener direction within each conversation, neural signals were copula-normalized separately for each participant. Then, the mutual information (GCMI) between these normalized signals across conversation partners was estimated for a range of speaker-listener time lags (-24 to 300 ms, in 4-ms steps) for each trial and frequency band. This process yielded a 4-D matrix per frequency band: trials × lags × speaker channels × listener channels.

From these matrices, we derived two complementary neural coordination measures: *concurrent* and *recurrent* inter-brain relationships (Burns et al., 2025) for each conversation trial. *Concurrent* neural coordination was defined as the peak speaker-listener GCMI within a narrow window around zero lag (-24 to 24 ms), capturing near-instantaneous coupling, whereas *recurrent* neural coordination was defined as the peak listener-trailing GCMI from 50 to 300 ms lags, corresponding to the canonical time window for speech processing. This approach allowed us to dissociate simultaneous from delayed forms of neural coordination during live conversation. The resulting GCMIIB values were then square-root transformed to improve normality (verified via Q-Q plot inspection) and standardized (z-scored).

We then asked whether the perceived quality of interaction predicts the strength of neural coordination across frequency bands. To test this, we fitted an LMM of GCMIIB as a function of PIQ for both concurrent and recurrent neural coordination at each frequency band. For each model, the fixed-effects structure included the PIQ × ROI-pair interaction, where ROI-pair comprised 36 levels corresponding to all pairwise combinations among the six ROIs per participant. Random intercepts for dyad, speaker (A or B) nested within dyad, and electrode-pair were included, as well as by-speaker random slopes for PIQ:

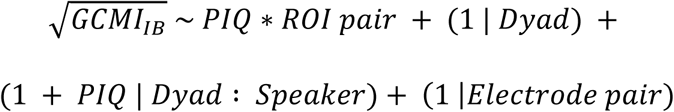

In models where the PIQ × ROI-pair interaction proved significant, we conducted follow-up EMT analyses to identify specific ROI pairs in which PIQ significantly predicted neural coordination (against zero). These 36 tests were corrected for multiple comparisons using the False Discovery Rate (FDR) method. We report these follow-up results only when EMT tests survive FDR correction.

In addition, to determine that our neural coordination results reflected genuine interpersonal coordination rather than spurious dependencies, we conducted a control analysis using pseudo-dyads (De Felice et al., 2024; Pan et al., 2018). Specifically, for each effect that was statistically significant and warranted interpretation in the main analysis, we generated 200 surrogate datasets by randomly shuffling dyad pairings while preserving the original sample size (24 dyads), number of trials, EEG channels, and data preparation steps. For each surrogate dataset, we fit the identical LMM used for the real dyads and extracted the corresponding test statistics to form a null distribution against which the observed effects were evaluated (Huang et al., 2025).

## 3. RESULTS

### 3.1. Speech-to-Speech (Prosodic Adaptation)

#### 3.1.1. Conversation-Level Synchrony

The prosodic synchrony model assessed whether a speaker’s prosodic features during a given turn were systematically influenced by their partner’s preceding turn. The model revealed a significant main effect of the partner’s preceding prosodic values (𝐹[1,25.88] = 89.44, 𝑝 < .001), indicating an overall synchrony effect. A significant interaction between the partner’s vocal expression and feature indicates that synchrony strength varied across acoustic features. Planned EMT analyses reveal that conversation partners tended to align their vocal modulations in F0 (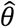 = 0.08, 95% CI [0.05,0.11], 𝑡[134] = 4.94, 𝑝 < .001), HNR (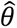 = 0.08, 95% CI [0.05,0.11], 𝑡[132] = 5.11, 𝑝 < .001), and especially loudness (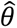 = 0.22, 95% CI [0.19,0.25], 𝑡[135] = 13.64, 𝑝 < .001). Voicing rate did not exhibit a significant slope (𝑝 = .105). We found no evidence that PIQ modulated these effects (*p*’s > .649).

#### 3.1.2. Conversation-Level Convergence

At the conversation level, there was a significant main effect of time (late – early) on convergence (𝐹[1,30.72] = 9.72, 𝑝 = .004), indicating that dyads became increasingly similar across acoustic features as conversations unfolded (see Figure 4, lower panel). Planned comparisons of model-derived EMTs indicated significant time slopes for F0 (𝛥*M* = 0.21, 95% CI [0.06, 0.36], *t*[267.98] = 2.70, *p* = .007) and voicing rate (𝛥*M* = 0.23, 95% CI [0.08, 0.38], *t*[267.98] = 3.07, *p* = .002), but not loudness or HNR (*p*’s > .314). However, the model yielded no evidence that PIQ modulated prosodic convergence at the conversation level (*p*’s > .364).

**Figure 4:**
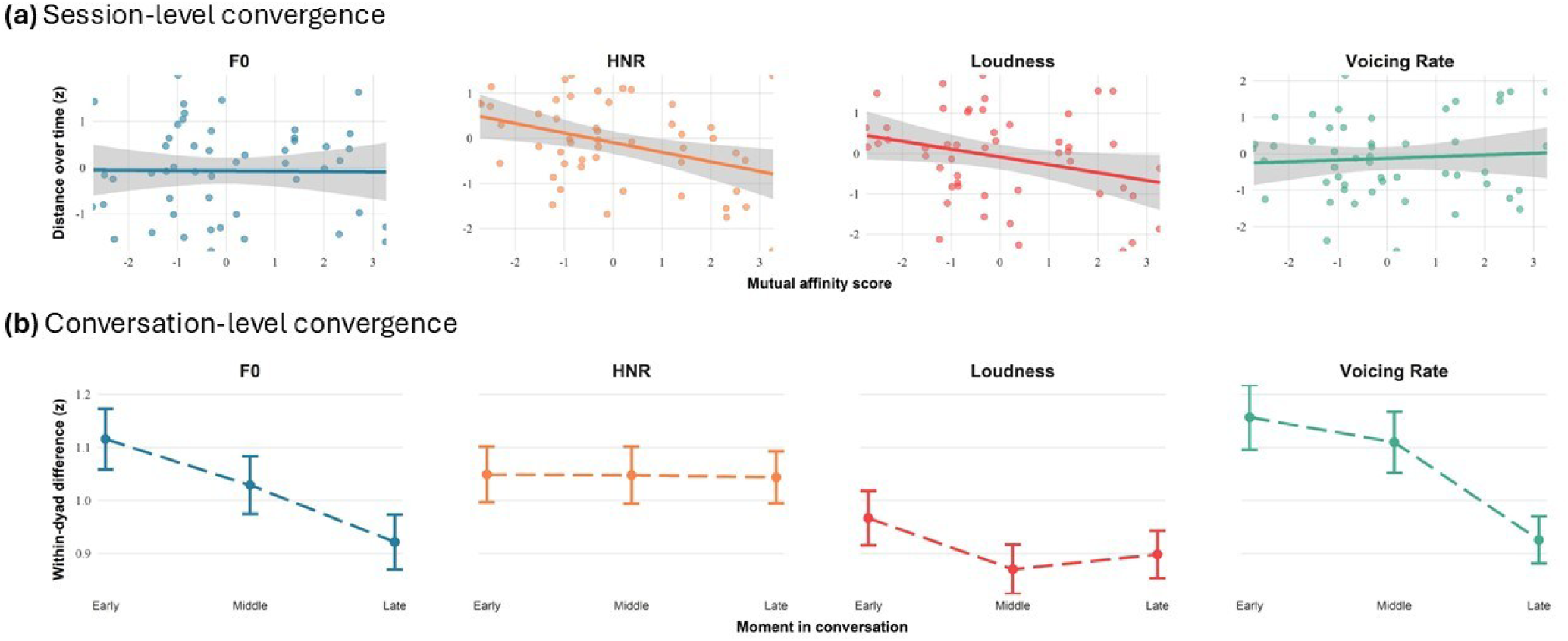
Prosodic convergence results. (a) Session-level convergence across acoustic features, showing the relationship between mutual affinity and partners’ decreasing acoustic distance. (b) Conversation-level convergence, shown by reductions in mean within-dyad differences from early to late moments in the interaction (Early: 0–80 s; Middle: 80–160 s; Late: 160–240 s). Shadowed areas and error bars represent SE.

#### 3.1.3. Session-Level Convergence

The session-level prosodic convergence model revealed a significant three-way interaction of Time × Feature × Mutual Affinity (𝐹[3,375.22] = 2.84, 𝑝 = .038), indicating that the association between mutual affinity and convergence over time differed across prosodic features (see Figure 4, upper panel). We conducted planned EMT analyses examining whether the slope of mutual affinity across time differed for each of the four theoretically motivated acoustic features. Results show that mutual affinity enhanced session-level convergence for HNR (𝛥*M* = 0.21, 95% CI [0.05, 0.37], *t*[187.29] = 2.62, *p* = .010) and loudness (𝛥*M* = 0.19, 95% CI [0.03, 0.36], *t*[187.29] = 2.38, *p* = .018), but not for F0 and voicing rate (*p*’s > .572).

### 3.2. Brain-to-Speech (Neural Speech Tracking)

The LMM for neural speech tracking, adjusting for acoustic covariates, yielded a significant interaction between PIQ and ROI (𝐹[5,11911.70] = 2.36, 𝑝 = .038), suggesting that the impact of PIQ on listeners’ neural tracking of their partner’s speech differs across cortical regions. In particular, we observed that the slope relating PIQ to neural speech tracking was significantly more positive in medial-central channels than in the rest of the ROIs (𝛥𝑀 = 0.05, 95% CI_Bonferroni(6)_ [0.00,0.09], 𝑡[11911.70] = 2.91, 𝑝_fdr[6]_ = .022). However, EMTs showed that the absolute slope in this medial-central ROI did not significantly differ from zero (𝜃^ = 0.08, 95% CI_Bonferroni(6)_ [−0.01,0.16], 𝑡[72.94] = 2.39, 𝑝_fdr[6]_ = .118). This indicates that PIQ modulated the topography of NST without producing a statistically reliable increase or decrease in the absolute level of tracking within any single ROI, controlling for conversation-level vocal-acoustic characteristics. Appendix B further explores these data and shows that the observed relative effect may be primarily driven by a single medial-central electrode rather than a broader regional pattern.

### 3.3. Brain-to-Brain (Neural Coordination / Inter-Brain Relationship)

#### 3.3.1. Theta Band

To test whether perceived interaction quality (PIQ) predicted theta-band neural coordination, we fitted LMMs that separately examined concurrent and recurrent coupling. The LMM for theta-band *concurrent* inter-brain relationship showed a significant interaction of PIQ and ROI pair (𝐹[35,251273.07] = 5.82, 𝑝 < .001). Follow-up EMTs revealed a significant effect of PIQ specifically at the medial-central to right-anterior ROI pair (𝜃^ = 0.06, 95% CI_Bonferroni[36]_ [0.00,0.12], 𝑡[103.83] = 3.44, 𝑝_fdr[36]_ = .031) and marginally non-significant effects at other medial-to-right ROI pairs: medial-anterior to right-anterior, medial-anterior to right-temporal, and medial-central to right-temporal pairs (*p*’s = .052). The LMM for theta-band *recurrent* neural coordination also exhibited a significant PIQ × ROI pair interaction (𝐹[35,251273.52] = 4.56, 𝑝 < .001), with follow-up EMTs indicating a significant relationship between PIQ and neural coordination only at the right-temporal to medial-anterior ROI pair (𝜃^ = 0.06, 95% CI_Bonferroni[36]_ [0.00,0.12], 𝑡[118.13] = 3.40, 𝑝_fdr[36]_ = .033. Appendix C displays the full EMT results. These results suggest that the perceived quality of social interaction is indexed, to some extent, by both concurrent and recurrent speaker-listener theta-band neural coordination, with effects localized to medial and right-lateralized regions.

#### 3.3.2. Alpha Band

We next fitted LMMs to test whether PIQ predicted alpha-band neural coordination across concurrent and recurrent (listener-trailing) time windows. The LMM for alpha-band *concurrent* neural coordination revealed a significant main effect of PIQ (𝐹[1,47.24] = 10.02, 𝑝 = .003), as well as of its interaction with ROI pair (𝐹[35,251272.57] = 5.09, 𝑝 < .001). Follow-up EMTs showed that higher PIQ was associated with greater alpha-band concurrent neural coordination at sixteen ROI pairs, primarily from speakers’ anterior ROIs (Table S3 in Appendix C; Figure 5). The strongest effects were observed for right-anterior to right-anterior (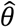 = 0.10, 95% CI_Bonferroni[36]_ [0.04,0.16], 𝑡[274.10] = 5.24, 𝑝_fdr[36]_ < .001), left-anterior to right-anterior (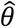 = 0.09, 95% CI_Bonferroni[36]_ [0.03,0.15], 𝑡[274.10] = 4.76, 𝑝_fdr[36]_ < .001), and right-anterior to medial-anterior (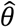 = 0.05, 95% CI_Bonferroni[36]_ [−0.01,0.10], 𝑡[198.29] = 2.87, 𝑝_fdr[36]_ = .015) ROI pairs.

**Figure 5:**
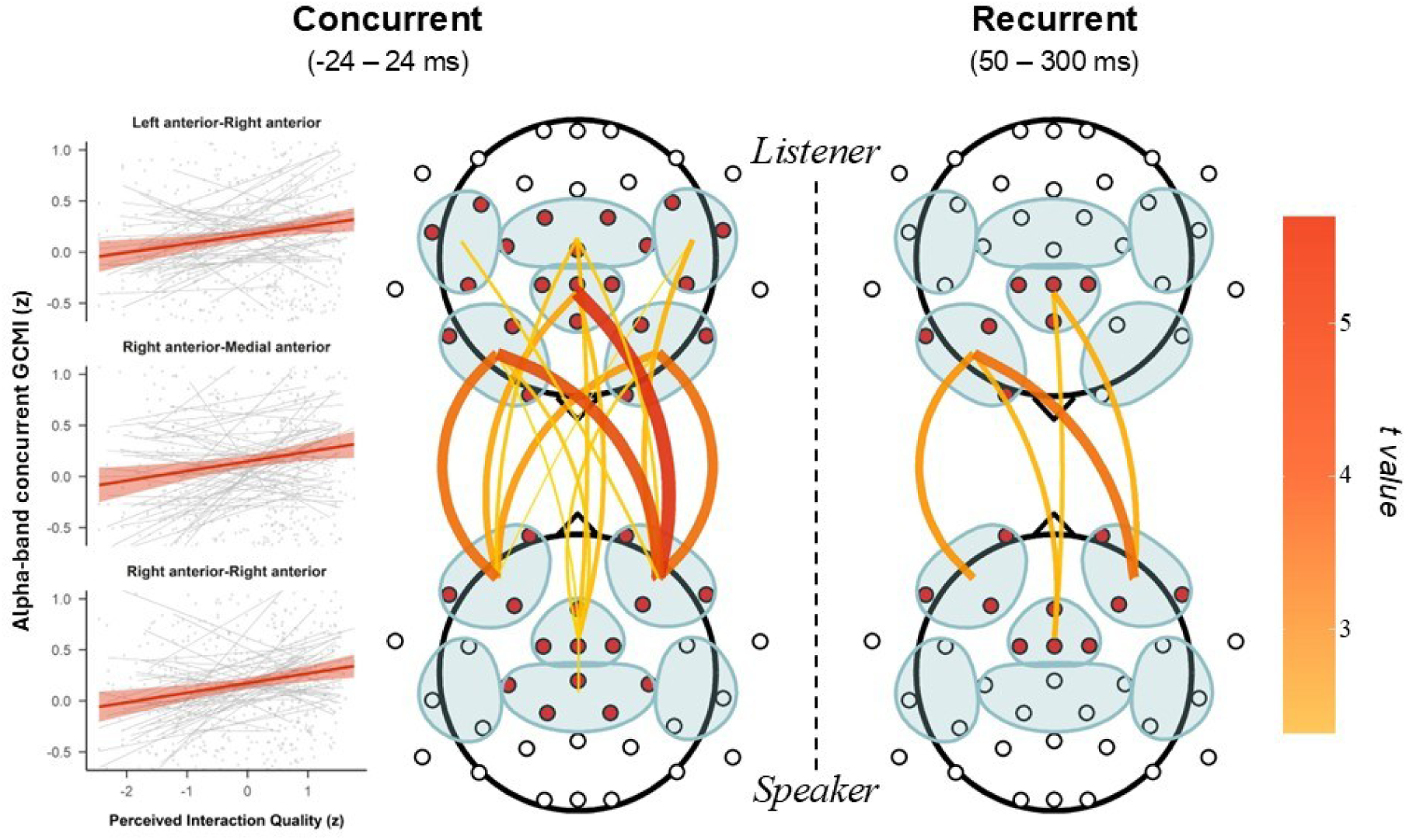
Alpha-band inter-brain relationships as a function of Perceived Interaction Quality (PIQ). Left: Concurrent links (–24 to 24 ms) for which the EMT simple slopes were significant. For three of these, we also show regression fits to the raw data, with thin gray lines illustrating dyad-level variability. Right: Recurrent links (50-300 ms) showing significant EMT simple slopes. In both panels, line color and thickness reflect the magnitude of the corresponding EMT simple-slope *t* value.

The LMM for alpha-band *recurrent* inter-brain relationship revealed a significant interaction between PIQ and ROI pair (𝐹[35,251273.77] = 4.16, 𝑝 < .001). Follow-up EMTs showed significant effects at five anterior ROI pairs: right-anterior to right-anterior (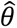 = 0.09, 95% CI_Bonferroni[36]_ [0.03,0.15], 𝑡[226.92] = 4.71, 𝑝_fdr[36]_ < .001), left-anterior to right- anterior (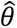 = 0.08, 95% CI_Bonferroni[36]_ [0.02,0.14], 𝑡[226.92] = 4.14, 𝑝_fdr[36]_ < .001), right-anterior to medial-anterior (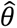 = 0.06, 95% CI_Bonferroni[36]_ [0.01,0.12], 𝑡[168.65] = 3.54, 𝑝_fdr[36]_ = .006), medial-anterior to right-anterior (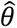 = 0.09, 95% CI_Bonferroni[36]_ [0.03,0.15], 𝑡[226.92] = 4.71, 𝑝_fdr[36]_ < .001), and medial-anterior to medial-anterior (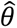 = 0.09, 95% CI_Bonferroni[36]_ [0.03,0.15], 𝑡[226.92] = 4.71, 𝑝_fdr[36]_ < .001). Together, these results suggest that higher interaction quality is associated with greater alpha-band neural coordination between conversation partners, with stronger effects for concurrent than recurrent coupling (Figure 5). Appendix D presents an exploratory analysis testing the association between neural coordination and mutual affinity scores.

#### 3.3.3. Control Analysis

Permutation tests confirmed that the estimated effects were specific to within-dyad interactions, as they were not observed in pseudo-dyads. Specifically, control analyses revealed that, for real dyads, the *F* statistic associated with the main effect of PIQ on alpha-band concurrent neural coordination exceeded that observed in all 200 surrogate samples (*p* = 0.005). The EMT simple slopes indexing the effects of PIQ on recurrent neural coordination at five frontal ROI pairs were also larger for real dyads than for any of the surrogate samples (*p* = 0.005). In the theta band, the observed EMT effects for both concurrent and recurrent inter-brain relationships exceeded most values obtained from pseudo-dyads (*p* = 0.02 and *p* = 0.01, respectively; see Figure 6).

**Figure 6:**
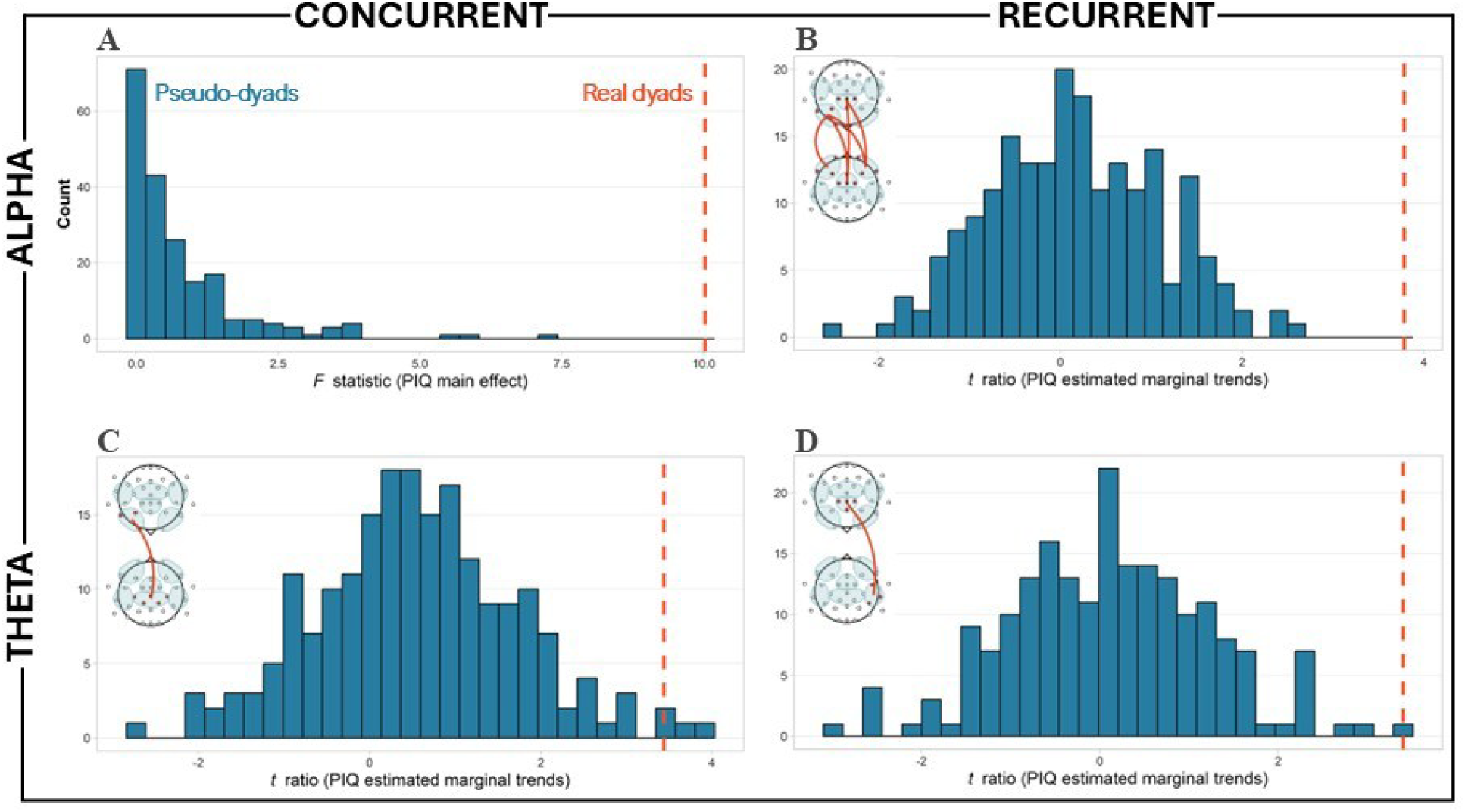
Null distributions from permutation control analyses comparing real dyads to pseudo-dyads. Null distributions (blue) were generated from 200 pseudo-dyad sample permutations. Orange dashed lines indicate statistics observed for real dyads. (A) Main effect of PIQ on concurrent alpha-band neural coordination (*F* statistic). (B) EMT *t* ratios for recurrent alpha coupling. Each observation is the average *t* ratio across all five significant ROI pairs. (C-D) EMT *t* ratios for concurrent theta coupling at medial-central to right-anterior channels and for recurrent theta coupling at right-temporal to medial-anterior channels, respectively.

## 4. DISCUSSION

The present study examined prosodic and neural markers of interpersonal alignment as a function of the jointly perceived quality of conversations and mutual affinity of conversational partners. At the *speech-to-speech* level, while dyads showed broadly generalized prosodic adaptation within individual conversations, mutual affinity between interlocutors modulated prosodic adaptation at the slower, session-level timescale. That is, over the course of the experiment, partners became increasingly similar in their vocal loudness and quality (HNR) when they reported more positive interpersonal impressions of one another. At the *brain-to-speech* level, we did not find reliable evidence that listeners’ neural tracking of their interlocutor’s speech varied in absolute magnitude as a function of perceived interaction quality (PIQ) within any region. At the *brain-to-brain* level, our main finding was that conversations perceived as higher in quality showed stronger inter-brain relationships across frequency bands (theta and alpha), with the most robust effects observed for concurrent (i.e., near-instantaneous) alpha-band neural coordination. Taken together, our findings highlight slow prosodic convergence as a marker of *relational* quality (i.e., mutual affinity between partners), and neural coordination—particularly in the alpha frequency band—as a robust marker of *conversational* quality (i.e., PIQ).

### 4.1. Mutual Affinity Enhances Slow Prosodic Convergence

It is well established that, as conversation unfolds, interlocutors tend to reciprocally adapt their vocal expression (Giles et al., 1991). Prosodic adaptation has been observed at various timescales: interlocutors may synchronize their turn-by-turn acoustic modulations (turn-level synchrony), converge over the course of a short interaction (conversation-level convergence), and/or converge slower, over the course of the whole experimental session (session-level convergence). Here, we assessed prosodic adaptation across these three timescales (see also Levitan & Hirschberg, 2011). Our results show that interlocutors tended to synchronize their turn-level prosodic modulations in loudness, F0 (pitch), and HNR (voice clarity), and significantly converged over the course of individual conversations in F0 and voicing rate. However, we did not find evidence that these effects were modulated by the perceived quality of the conversation. It is noteworthy that while we found PIQ to robustly modulate the extent of neural coordination (see below), we did not observe comparable evidence at the speech-to-speech level. The interactive-alignment model makes the prediction that interpersonal alignment operates across levels, and “percolates” between levels (Menenti et al., 2012). Accordingly, we expected alignment associated with PIQ to percolate from neural dynamics to overt vocal behavior. Instead, our results echo the argument in Ostrand & Chodroff (2021) that alignment is not a unitary phenomenon; instead, interpersonal alignment at different levels may be sensitive to distinct features of the communicative situation. Accordingly, they predict that alignment can occur for some levels, and not occur for others.

Over the course of the experimental session (∼ 1.5 h), however, newly acquainted individuals exhibited systematically stronger prosodic convergence as a function of *mutual affinity*: partners who liked and trusted one another more displayed more similar volume and voice quality (HNR) over time. This is in accordance with our hypothesis and prior evidence. For instance, Pardo et al. (2012) followed pairs of roommates over the course of a school year and found that their degree of phonetic convergence (i.e., the extent to which their pronunciations of a predefined set of words became more similar) was related to how close they reported feeling to one another after 3.5 months. Although that study spans a longer time scale and focuses on phonetic rather than prosodic features, both sets of findings are consistent with predictions derived from Communication Accommodation Theory (Giles et al., 1973, 1991). This theory proposes that accommodation phenomena—such as prosodic and phonetic adaptation—both index and seek to establish solidarity (or dissociation) with an interaction partner. Specifically, by speaking more similarly, which typically happens unconsciously, interlocutors reduce interpersonal differences and foster social integration (Giles et al., 1991). Our results are consistent with the view that prosodic convergence is a relationship-sensitive process that reflects, and likely contributes to, social affiliation.

### 4.2. Perceived Interaction Quality Modulates Neural Coordination

In this study, naturalistic variation in conversational quality was promoted by prompting newly acquainted pairs to discuss topics that were either highly engaging or minimally engaging for both partners. Importantly, however, our primary index of *perceived* interaction quality (i.e., PIQ) was a composite behavioral index integrating reward value (in the form of time bidding), perceived effort, and topic interest, thereby capturing how participants evaluated each interaction across multiple psychologically meaningful dimensions. Our analyses demonstrate a positive association between PIQ and neural coordination between conversation partners, with the most robust effects emerging for concurrent alpha-band coupling, predominantly involving anterior electrode sites. These findings are in line with previous studies of verbal interaction that have reliably observed concurrent neural coordination within prefrontal and temporo-parietal regions, presumably reflecting their involvement in social cognition, mutual attention, and the language network (Gvirts & Perlmutter, 2020; Kelsen et al., 2022; Pérez et al., 2019; Schwartz et al., 2024).

For instance, one study using EEG hyperscanning (Pérez et al., 2019) found that conversation in a foreign language was associated with a distinct and more widespread pattern of alpha-band (but not theta- or beta-band) neural coordination over fronto-central, temporal, and parietal regions relative to native-language conversation; these results were interpreted as reflecting a shift in joint attentional strategies in response to the cognitive demands imposed by different linguistic contexts. Another study (Pan et al., 2018) employed fNIRS hyperscanning while a trained instructor taught a song to a learner using two instructional formats: a low-interactive approach that prioritized uninterrupted exposure to the material, and a highly interactive approach involving frequent turn-taking and self-management of the interaction. They found that the high-interactive condition resulted in significantly higher inter-brain relationships in the bilateral inferior frontal cortex, which in turn was positively associated with learning outcomes (solo song performance). The authors suggested that neural coordination in these regions reflects a neural marker of active information transfer facilitated by *high-involvement* interactions. Together with these studies, our results are in accordance with the view that perceived high-quality, *engaging* conversations are characterized by the deployment of shared-attentional strategies and effective information transfer between interlocutors (De Felice et al., 2025; Hasson & Frith, 2016; Liu et al., 2021), possibly supported by shared representations of the communicative situation (Hasson et al., 2012; Menenti et al., 2012; Pickering & Garrod, 2004; Stolk et al., 2016).

The neurofunctional model of interpersonal alignment proposed in Shamay-Tsoory et al. (2019) and elaborated in Gvirts & Perlmutter (2020) conceptualizes mutual attention as a foundational mechanism of successful interactions. This framework proposes that socially *significant* interactions—i.e., those carrying high reward value—facilitate shared attentional engagement through the coupling of interlocutors’ *mutual social attention systems*, encompassing prefrontal and/or temporoparietal brain regions (Gvirts & Perlmutter, 2020). These shared attentional processes are thought to optimize cognitive processing and to be reinforced via the recruitment of reward-related neural systems, particularly when interlocutors successfully coordinate at behavioral, emotional, and cognitive levels (Shamay-Tsoory et al., 2019).

Moreover, frontal alpha-band neural oscillations have been robustly associated with attentional processing and predictive top-down control (Dikker et al., 2017; Kayser et al., 2015; Park et al., 2015). Thus, the dominant alpha-band, concurrent inter-brain relationship in our data is consistent with the model of *interaction-level* interpersonal alignment and suggests that conversations jointly perceived as engaging, effortless, and focused on topics of mutual interest are more likely to promote mutual engagement of social-attentional neural systems.

Notably, our data also show robust recurrent (i.e., listener-trailing) coupling in the alpha band. This pattern implies a temporal delay on the order of speech production and subsequent perceptual processing, suggesting coupling mediated by the speech signal itself (Pérez et al., 2022). That is, recurrent coupling may be interpreted as indexing *perceptual-level* alignment facilitated by sequential processing along the speaker-listener chain (Denes & Pinson, 1993).

However, given the spatial overlap between recurrent and concurrent effects (see Figure 5), an alternative interpretation is conceivable: the observed pattern of recurrent coordination may reflect sustained attentional engagement with the speaker rather than a distinct perceptual process. Specifically, higher PIQ may promote sustained listener engagement at any given point, producing strong concurrent coordination that carries over into short-lag dependencies through continuous, speaker-driven updating of the listener’s neural state. Consistent with this interpretation, PIQ did not robustly modulate neural speech tracking, a sensitive marker of perceptual encoding, suggesting that the observed effects are unlikely to reflect enhanced processing of the speech signal.

Although less robust, we also observed an effect of PIQ on neural coordination in the theta band, involving medial-central to right-anterior concurrent coupling and right-temporal to medial-anterior recurrent coupling. We had hypothesized stronger theta coordination given this frequency’s key role in behavioral coordination (Wang et al., 2020) as well as speech segmentation and paralinguistic processing (Poeppel & Assaneo, 2020). Specifically, in the fully free-form conversational format of our paradigm, we argue that conversations perceived as high in quality are likely characterized by smoother joint action coordination, possibly driving the increased *concurrent* theta coupling (M. Yang et al., 2023). *Recurrent* theta coupling, on the other hand, may reflect upregulated listener processing of the (para)linguistic signal, resulting in greater speaker-listener similarity along the speech chain (Pérez et al., 2022). While we did not find reliable modulation of online speech encoding (i.e., neural speech tracking) by PIQ in the listener to meaningfully support this hypothesis, it exhibited relative effects localized to medial-central channels (see also Appendix B). These findings neither confirm nor rule out the possibility that variation in interaction quality modulates speech processing in a way that facilitates the recurrent inter-brain relationships observed here.

In summary, the present findings show, for the first time, that the perceived quality of a social interaction is reflected in the extent of interpersonal neural coordination across frequency bands and temporal dependencies, spanning concurrent and listener-lagging coordination.

However, the predominance of alpha-band concurrent inter-brain relationships suggests that a *good* conversation is primarily characterized by an enhancement of interaction-level (rather than sensory/perceptual) processes, such as the co-construction of situation models or common ground (Menenti et al., 2012), convergence in affective or motivational states (Hoehl et al., 2021), or an overall state of optimal social integration (Burns et al., 2025).

### 4.3. Strengths and Limitations

A limitation of this and similar paradigms is that their correlational design precludes inferences about causality (Novembre & Iannetti, 2021). The quality of each conversation, shaped by its topic and the interlocutors’ mutual affinity, likely influences interpersonal alignment and coordination, but more coordinated interactions may also influence the subjective experience of the conversation and the interlocutors’ relationship. The relationship is likely bidirectional, with interpersonal alignment both reflecting and contributing to interaction quality. On the one hand, conversations about a topic we are passionate about may hold greater reward value, leading to heightened and more coordinated engagement of the mutual social attention system (Gvirts & Perlmutter, 2020; Shamay-Tsoory et al., 2019). On the other hand, accumulating evidence indicates that neural coordination may itself exert a causal influence on interpersonal outcomes (Leiva-Cisterna et al., 2025; Novembre et al., 2017; Novembre & Iannetti, 2021; Pan et al., 2021; Y. Yang et al., 2021), and that behavioral coordination serves as a potent relational signal to foster perceived positive interactions (Cohen et al., 2024; Hoehl et al., 2021; Kokal et al., 2011).

An important consideration when interpreting the present findings is that all dyads consisted of newly acquainted individuals who had never met before and did not get to interact until the start of the experiment. Generalization to established relationships (e.g., friends or romantic partners) may be limited. Converging evidence suggests that strangers are more likely to exhibit greater alignment than partners with an established connection in naturalistic interaction. For example, in an empathy-giving task using EEG hyperscanning, couples showed lower alpha-band neural coordination despite higher behavioral coordination, whereas strangers showed the opposite pattern. This was taken to suggest that unfamiliar partners must recruit greater neural alignment through effortful attention to support empathy, whereas long-term partners can rely on efficient, automatic coordination through well-practiced interpersonal routines (Djalovski et al., 2021). In a study using fMRI hyperscanning and natural language processing to track the neural and linguistic trajectories of friends and strangers during conversation, strangers tended to converge by becoming more neurally and linguistically similar to establish common ground, whereas friends tended to diverge by exploring a broader range of novel ideas and neural states. Interestingly, when strangers adopted a more exploratory behaviour (like friends did), they reported higher levels of enjoyment and closeness with their partner (Speer et al., 2024). Indeed, more interpersonal alignment is not always better; engaging social interactions appear to depend on a balance between predictability, which supports mutual alignment, and novelty/complexity (Ravreby et al., 2022). Together, these findings suggest that among partners with an established social bond, optimal social interaction relies on flexible modulation of alignment, whereas unacquainted dyads may prioritize alignment as much as possible as a strategy for establishing common ground and foster conversational quality. Future research should explicitly address this hypothesis.

An important strength of the present study lies in its use of a minimally constrained conversational paradigm, allowing interlocutors to self-manage their interaction dynamics, engaging in naturalistic turn-taking and backchanneling in a way that resembles everyday interaction. It is under these conditions that interlocutors must anticipate turn completions, plan appropriate responses, and monitor their partner’s reactions in real time (Templeton et al., 2022), making such a paradigm particularly valuable for studies on interpersonal alignment and coordination. In addition, this work responds to recent calls to incorporate information-theoretic metrics in hyperscanning research (Chidichimo et al., 2025), showing that the Gaussian Copula Mutual Information framework (Ince et al., 2017) provides a useful index of cross-brain signal dependencies in EEG hyperscanning. Future research should extend these approaches to longitudinal designs to determine whether short-term alignment during early social interactions can predict downstream outcomes in social affiliation (e.g., Shen et al., 2025).

### 4.4. Conclusion

In sum, by jointly measuring speech-to-speech, brain-to-speech, and brain-to-brain dependencies between partners during naturalistic conversation, the present study provides an integrated account of how the quality of interaction and interpersonal affinity are instantiated in shared behavioral (i.e., vocal) and neural dynamics at different timescales. Our findings characterize interpersonal alignment as a multilevel phenomenon that unfolds across functional domains, each indexing distinct aspects of social interaction. Slow prosodic convergence in interlocutors’ loudness and vocal clarity reflected the gradual formation of their relationship, increasing across repeated conversations as partners developed more positive interpersonal impressions. By contrast, inter-brain relationships across frequency bands (theta and alpha) and temporal dependencies (concurrent and recurrent) tracked the quality of individual conversations, such that interactions experienced as interesting, effortless, and worth sustained engagement were marked by stronger neural coordination, most prominently for concurrent alpha-band coupling. This dissociation suggests that interpersonal alignment is not a unitary phenomenon, but a multilevel process through which humans simultaneously regulate proximal (ongoing interaction experience) and distal (relationship formation) social outcomes. These findings provide novel insights into *how* and *when* alignment mechanisms map onto distinct social outcomes and extend emerging perspectives in the growing field of Relational Neuroscience (De Felice et al., 2025).

## Acknowledgements

We are sincerely grateful to Daphne Yu for her valuable assistance with data (pre)processing. We also thank Dr. Karsten Steinhauer for valuable feedback during the early stages of this project.

This work was supported by a Discovery Grant (RGPIN-2022-04363) awarded to M.D.P and a fellowship from the Secretary of Science, Humanities and Innovation (SECIHTI) of Mexico to M.E.D. (award number: 834007). A.P. was supported by the Agencia Estatal de Investigación (AEI), Spain [Grant PID2024-157011NB-I00], co-funded by the European Union.

## Data and code availability statement

All (pre)processing and analysis code required to reproduce the results is publicly available at https://github.com/elidom/EEG-Hyperscanning-Code_How-do-we-align-in-good-conversations-project. The preprocessed datasets necessary to reproduce the reported analyses will also be made publicly available upon acceptance of the manuscript for publication.

