## Appendix A for "How do we align in good conversation? Investigating the link between interaction quality and multimodal interpersonal coordination"

**Appendix A.** Mutual affinity questionnaire. Adapted from Montepare et al. (2014).

Please answer the following questions about your partner. Your responses are completely anonymous and confidential. Your honest answers will help us obtain more accurate results.

| | 1 = Strongly Disagree | | | / | | | 7 = Strongly Agree |
|---|---|---|---|---|---|---|---|
| This person is approachable | 1 | 2 | 3 | 4 | 5 | 6 | 7 |
| This person is competent | 1 | 2 | 3 | 4 | 5 | 6 | 7 |
| I would like to become friends with this person | 1 | 2 | 3 | 4 | 5 | 6 | 7 |
| This person would make me feel at ease | 1 | 2 | 3 | 4 | 5 | 6 | 7 |
| This person would be difficult to talk to | 1 | 2 | 3 | 4 | 5 | 6 | 7 |
| This person is attractive | 1 | 2 | 3 | 4 | 5 | 6 | 7 |
| This person is a poor listener | 1 | 2 | 3 | 4 | 5 | 6 | 7 |
| I would like to work with this person | 1 | 2 | 3 | 4 | 5 | 6 | 7 |
| This person is friendly | 1 | 2 | 3 | 4 | 5 | 6 | 7 |
| I would avoid this person | 1 | 2 | 3 | 4 | 5 | 6 | 7 |
| This person would be a good storyteller | 1 | 2 | 3 | 4 | 5 | 6 | 7 |
| This person is trustworthy | 1 | 2 | 3 | 4 | 5 | 6 | 7 |
