## Appendix B for "How do we align in good conversation? Investigating the link between interaction quality and multimodal interpersonal coordination"

**Appendix B: Neural speech tracking post-hoc examination**

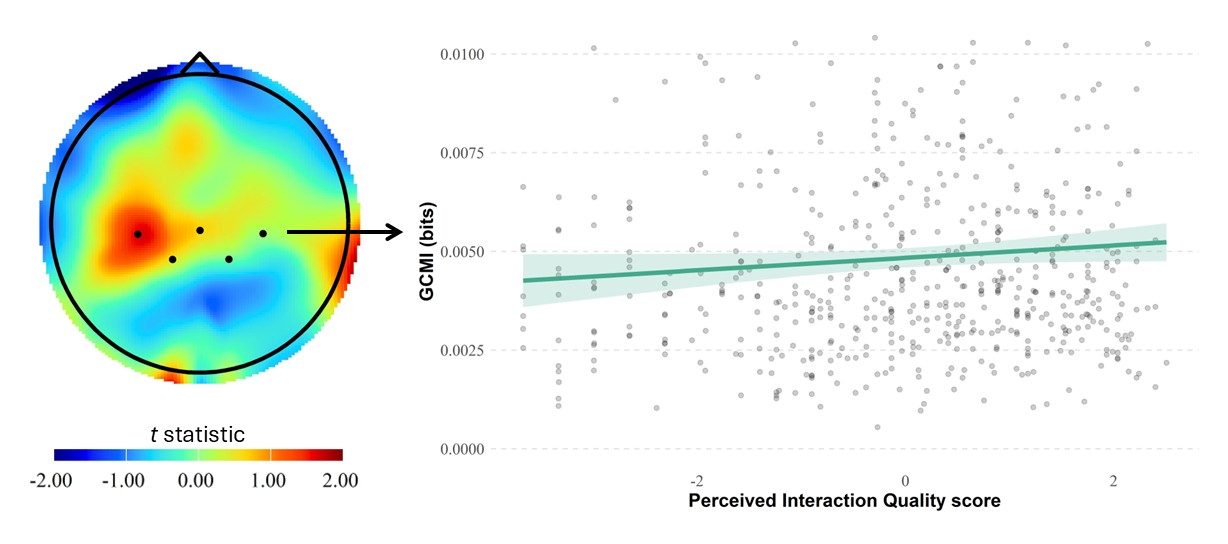
In section 3.2. of the main text, we did not find reliable effects of perceived interaction quality (PIQ) on neural speech tracking (NST). Specifically, the PIQ × ROI interaction was significant in the overall model; however, follow-up estimated marginal trend (EMT) analyses revealed no ROI-specific simple slope significantly different from zero after FDR correction for six ROIs. Note, however, that a positive slope at medial-central channels was present at the *uncorrected* level. Exploratory data analysis (Figure S1) suggests that the association between PIQ and NST is concentrated around *one* medial central electrode (C3). Thus, it is likely that the PIQ-NST relationship, if real, is confined to this focal region, making it difficult to detect given our sensor density and pre-defined ROI framework. Future work testing similar hypotheses would benefit from denser sensor arrays and data-driven, cluster-based inference approaches.

**Figure S1:** Neural speech tracking results. Left: Scalp topography showing the t-statistics from channel-wise linear regressions of GCMI on PIQ scores. The black dots indicate the placement of medial-central electrodes. Right: Regression plot showing the relationship between perceived interaction quality and medial-central GCMI; shaded band indicates the SE. Simple-slope estimation via estimated marginal trends of our LMM showed that this slope did not differ significantly from 0, controlling for acoustic covariates, and after FDR correction for multiple comparisons.

As an exploratory analysis, we examined whether individual affinity scores (from person-perception ratings) modulated the extent of NST—that is, whether greater liking of one’s interlocutor was associated with enhanced neural encoding of their speech. An LMM predicting GCMI_NST_ from affinity scores and their interaction with ROI (including random intercepts for dyad, listener-within-dyad, and trial) showed a marginal *negative* association with affinity ($F\left[ 1,45 \right]=3.62$, $p=.063$).
