## Appendix C for "How do we align in good conversation? Investigating the link between interaction quality and multimodal interpersonal coordination"

**Appendix C. Neural coordination estimated marginal trends (EMT) tables.**

**Table S1.** EMTs for the effect of PIQ on *theta-band concurrent* neural coordination

| ROI Pair | Estimate | Std. Error | df | Lower CI | Upper CI | p-value |
| --- | --- | --- | --- | --- | --- | --- |
| Left anterior-Left anterior | 0.00 | 0.02 | 155.40 | -0.06 | 0.07 | 0.912 |
| Left anterior-Left temporal | -0.01 | 0.02 | 155.40 | -0.07 | 0.05 | 0.711 |
| Left anterior-Medial anterior | 0.01 | 0.02 | 122.00 | -0.04 | 0.07 | 0.641 |
| Left anterior-Medial central | -0.03 | 0.02 | 103.80 | -0.08 | 0.03 | 0.358 |
| Left anterior-Right anterior | 0.01 | 0.02 | 155.40 | -0.05 | 0.07 | 0.711 |
| Left anterior-Right temporal | 0.02 | 0.02 | 155.40 | -0.04 | 0.08 | 0.457 |
| Left temporal-Left anterior | 0.03 | 0.02 | 155.40 | -0.03 | 0.09 | 0.358 |
| Left temporal-Left temporal | 0.01 | 0.02 | 155.40 | -0.06 | 0.07 | 0.87 |
| Left temporal-Medial anterior | 0.00 | 0.02 | 122.00 | -0.06 | 0.06 | 0.913 |
| Left temporal-Medial central | -0.01 | 0.02 | 103.80 | -0.07 | 0.05 | 0.711 |
| Left temporal-Right anterior | 0.05 | 0.02 | 155.40 | -0.01 | 0.11 | 0.07 |
| Left temporal-Right temporal | -0.00 | 0.02 | 155.40 | -0.06 | 0.06 | 0.92 |
| Medial anterior-Left anterior | 0.02 | 0.02 | 122.00 | -0.03 | 0.08 | 0.358 |
| Medial anterior-Left temporal | 0.03 | 0.02 | 122.00 | -0.03 | 0.09 | 0.358 |
| Medial anterior-Medial anterior | 0.02 | 0.02 | 99.50 | -0.03 | 0.08 | 0.358 |
| Medial anterior-Medial central | -0.00 | 0.02 | 87.10 | -0.06 | 0.05 | 0.913 |
| Medial anterior-Right anterior | 0.05 | 0.02 | 122.00 | -0.01 | 0.11 | 0.052 |
| Medial anterior-Right temporal | 0.05 | 0.02 | 122.00 | -0.01 | 0.11 | 0.052 |
| Medial central-Left anterior | 0.03 | 0.02 | 103.80 | -0.02 | 0.09 | 0.301 |
| Medial central-Left temporal | 0.02 | 0.02 | 103.80 | -0.03 | 0.08 | 0.358 |
| Medial central-Medial anterior | -0.01 | 0.02 | 87.10 | -0.07 | 0.04 | 0.602 |
| Medial central-Medial central | -0.03 | 0.02 | 77.80 | -0.09 | 0.02 | 0.265 |
| Medial central-Right anterior | 0.06 | 0.02 | 103.80 | 0.00 | 0.12 | 0.031* |
| Medial central-Right temporal | 0.05 | 0.02 | 103.80 | -0.01 | 0.11 | 0.052 |
| Right anterior-Left anterior | -0.00 | 0.02 | 155.40 | -0.07 | 0.06 | 0.904 |
| Right anterior-Left temporal | -0.02 | 0.02 | 155.40 | -0.08 | 0.04 | 0.586 |
| Right anterior-Medial anterior | 0.01 | 0.02 | 122.00 | -0.05 | 0.07 | 0.646 |
| Right anterior-Medial central | -0.03 | 0.02 | 103.80 | -0.09 | 0.02 | 0.276 |
| Right anterior-Right anterior | 0.01 | 0.02 | 155.40 | -0.06 | 0.07 | 0.868 |
| Right anterior-Right temporal | 0.03 | 0.02 | 155.40 | -0.03 | 0.09 | 0.358 |
| Right temporal-Left anterior | 0.03 | 0.02 | 155.40 | -0.03 | 0.09 | 0.358 |
| Right temporal-Left temporal | -0.01 | 0.02 | 155.40 | -0.08 | 0.05 | 0.646 |
| Right temporal-Medial anterior | 0.02 | 0.02 | 122.00 | -0.04 | 0.08 | 0.457 |
| Right temporal-Medial central | -0.02 | 0.02 | 103.80 | -0.08 | 0.04 | 0.388 |
| Right temporal-Right anterior | 0.03 | 0.02 | 155.40 | -0.04 | 0.09 | 0.358 |
| Right temporal-Right temporal | 0.02 | 0.02 | 155.40 | -0.04 | 0.09 | 0.388 |

Note. p-values are FDR-corrected.

**Table S2.** EMTs for the effect of PIQ on *theta-band recurrent* neural coordination

| ROI Pair | Estimate | Std. Error | df | Lower CI | Upper CI | p-value |
| --- | --- | --- | --- | --- | --- | --- |
| Left anterior-Left anterior | 0.03 | 0.02 | 149.50 | -0.04 | 0.09 | 0.631 |
| Left anterior-Left temporal | -0.01 | 0.02 | 149.50 | -0.08 | 0.05 | 0.792 |
| Left anterior-Medial anterior | 0.04 | 0.02 | 118.10 | -0.02 | 0.10 | 0.218 |
| Left anterior-Medial central | 0.02 | 0.02 | 101.10 | -0.04 | 0.07 | 0.671 |
| Left anterior-Right anterior | 0.02 | 0.02 | 149.50 | -0.04 | 0.08 | 0.671 |
| Left anterior-Right temporal | 0.02 | 0.02 | 149.50 | -0.04 | 0.08 | 0.671 |
| Left temporal-Left anterior | -0.00 | 0.02 | 149.50 | -0.07 | 0.06 | 0.878 |
| Left temporal-Left temporal | 0.00 | 0.02 | 149.50 | -0.06 | 0.07 | 0.921 |
| Left temporal-Medial anterior | 0.04 | 0.02 | 118.10 | -0.02 | 0.10 | 0.218 |
| Left temporal-Medial central | 0.01 | 0.02 | 101.10 | -0.04 | 0.07 | 0.692 |
| Left temporal-Right anterior | 0.01 | 0.02 | 149.50 | -0.06 | 0.07 | 0.877 |
| Left temporal-Right temporal | -0.02 | 0.02 | 149.50 | -0.09 | 0.04 | 0.631 |
| Medial anterior-Left anterior | 0.00 | 0.02 | 118.10 | -0.06 | 0.07 | 0.878 |
| Medial anterior-Left temporal | -0.03 | 0.02 | 118.10 | -0.09 | 0.03 | 0.551 |
| Medial anterior-Medial anterior | -0.00 | 0.02 | 97.00 | -0.06 | 0.05 | 0.907 |
| Medial anterior-Medial central | -0.02 | 0.02 | 85.30 | -0.07 | 0.04 | 0.671 |
| Medial anterior-Right anterior | -0.01 | 0.02 | 118.10 | -0.07 | 0.05 | 0.805 |
| Medial anterior-Right temporal | -0.00 | 0.02 | 118.10 | -0.06 | 0.06 | 0.907 |
| Medial central-Left anterior | -0.01 | 0.02 | 101.10 | -0.06 | 0.05 | 0.877 |
| Medial central-Left temporal | 0.01 | 0.02 | 101.10 | -0.05 | 0.06 | 0.877 |
| Medial central-Medial anterior | -0.04 | 0.02 | 85.30 | -0.10 | 0.02 | 0.218 |
| Medial central-Medial central | -0.02 | 0.02 | 76.50 | -0.08 | 0.03 | 0.551 |
| Medial central-Right anterior | -0.02 | 0.02 | 101.10 | -0.07 | 0.04 | 0.671 |
| Medial central-Right temporal | 0.01 | 0.02 | 101.10 | -0.04 | 0.07 | 0.671 |
| Right anterior-Left anterior | 0.04 | 0.02 | 149.50 | -0.03 | 0.10 | 0.386 |
| Right anterior-Left temporal | -0.01 | 0.02 | 149.50 | -0.07 | 0.05 | 0.877 |
| Right anterior-Medial anterior | 0.04 | 0.02 | 118.10 | -0.02 | 0.10 | 0.273 |
| Right anterior-Medial central | 0.02 | 0.02 | 101.10 | -0.04 | 0.07 | 0.671 |
| Right anterior-Right anterior | 0.02 | 0.02 | 149.50 | -0.05 | 0.08 | 0.671 |
| Right anterior-Right temporal | 0.01 | 0.02 | 149.50 | -0.06 | 0.07 | 0.877 |
| Right temporal-Left anterior | 0.02 | 0.02 | 149.50 | -0.04 | 0.08 | 0.671 |
| Right temporal-Left temporal | 0.00 | 0.02 | 149.50 | -0.06 | 0.06 | 0.976 |
| Right temporal-Medial anterior | 0.06 | 0.02 | 118.10 | 0.00 | 0.12 | 0.033* |
| Right temporal-Medial central | 0.02 | 0.02 | 101.10 | -0.04 | 0.08 | 0.651 |
| Right temporal-Right anterior | 0.03 | 0.02 | 149.50 | -0.03 | 0.09 | 0.551 |
| Right temporal-Right temporal | 0.02 | 0.02 | 149.50 | -0.04 | 0.09 | 0.631 |

Note. p-values are FDR-corrected.

**Table S3.** EMTs for the effect of PIQ on *alpha-band concurrent* neural coordination

| ROI Pair | Estimate | Std. Error | df | Lower CI | Upper CI | p-value |
| --- | --- | --- | --- | --- | --- | --- |
| Left anterior-Left anterior | 0.07 | 0.02 | 274.10 | 0.01 | 0.13 | <0.001*** |
| Left anterior-Left temporal | 0.04 | 0.02 | 274.10 | -0.02 | 0.10 | 0.0469* |
| Left anterior-Medial anterior | 0.06 | 0.02 | 198.30 | 0.01 | 0.12 | 0.0013** |
| Left anterior-Medial central | 0.05 | 0.02 | 158.70 | -0.00 | 0.10 | 0.0134* |
| Left anterior-Right anterior | 0.09 | 0.02 | 274.10 | 0.03 | 0.15 | <0.001*** |
| Left anterior-Right temporal | 0.04 | 0.02 | 274.10 | -0.02 | 0.10 | 0.0912 |
| Left temporal-Left anterior | 0.02 | 0.02 | 274.10 | -0.04 | 0.08 | 0.4737 |
| Left temporal-Left temporal | 0.03 | 0.02 | 274.10 | -0.03 | 0.09 | 0.1691 |
| Left temporal-Medial anterior | 0.02 | 0.02 | 198.30 | -0.03 | 0.08 | 0.2118 |
| Left temporal-Medial central | 0.01 | 0.02 | 158.70 | -0.05 | 0.06 | 0.7947 |
| Left temporal-Right anterior | 0.04 | 0.02 | 274.10 | -0.02 | 0.10 | 0.0751 |
| Left temporal-Right temporal | 0.04 | 0.02 | 274.10 | -0.02 | 0.10 | 0.0681 |
| Medial anterior-Left anterior | 0.05 | 0.02 | 198.30 | -0.01 | 0.10 | 0.0186* |
| Medial anterior-Left temporal | 0.02 | 0.02 | 198.30 | -0.03 | 0.08 | 0.2463 |
| Medial anterior-Medial anterior | 0.05 | 0.02 | 149.50 | -0.00 | 0.10 | 0.0085** |
| Medial anterior-Medial central | 0.05 | 0.01 | 123.50 | -0.00 | 0.10 | 0.0115* |
| Medial anterior-Right anterior | 0.05 | 0.02 | 198.30 | -0.01 | 0.10 | 0.0149* |
| Medial anterior-Right temporal | 0.02 | 0.02 | 198.30 | -0.03 | 0.08 | 0.2064 |
| Medial central-Left anterior | 0.02 | 0.02 | 158.70 | -0.04 | 0.07 | 0.3746 |
| Medial central-Left temporal | -0.00 | 0.02 | 158.70 | -0.06 | 0.05 | 0.8522 |
| Medial central-Medial anterior | 0.02 | 0.01 | 123.50 | -0.03 | 0.07 | 0.2064 |
| Medial central-Medial central | 0.00 | 0.01 | 104.40 | -0.04 | 0.05 | 0.857 |
| Medial central-Right anterior | 0.04 | 0.02 | 158.70 | -0.01 | 0.09 | 0.0315* |
| Medial central-Right temporal | -0.01 | 0.02 | 158.70 | -0.06 | 0.04 | 0.5059 |
| Right anterior-Left anterior | 0.09 | 0.02 | 274.10 | 0.03 | 0.15 | <0.001*** |
| Right anterior-Left temporal | 0.06 | 0.02 | 274.10 | 0.00 | 0.12 | 0.0046** |
| Right anterior-Medial anterior | 0.10 | 0.02 | 198.30 | 0.04 | 0.15 | <0.001*** |
| Right anterior-Medial central | 0.04 | 0.02 | 158.70 | -0.01 | 0.10 | 0.0159* |
| Right anterior-Right anterior | 0.10 | 0.02 | 274.10 | 0.04 | 0.16 | <0.001*** |
| Right anterior-Right temporal | 0.05 | 0.02 | 274.10 | -0.01 | 0.11 | 0.0186* |
| Right temporal-Left anterior | -0.00 | 0.02 | 274.10 | -0.06 | 0.06 | 0.95 |
| Right temporal-Left temporal | 0.03 | 0.02 | 274.10 | -0.03 | 0.09 | 0.117 |
| Right temporal-Medial anterior | 0.02 | 0.02 | 198.30 | -0.03 | 0.08 | 0.2064 |
| Right temporal-Medial central | 0.00 | 0.02 | 158.70 | -0.05 | 0.06 | 0.892 |
| Right temporal-Right anterior | 0.00 | 0.02 | 274.10 | -0.06 | 0.06 | 0.892 |
| Right temporal-Right temporal | 0.04 | 0.02 | 274.10 | -0.02 | 0.10 | 0.0629 |

Note. p-values are FDR-corrected.

**Table S4.** EMTs for the effect of PIQ on *alpha-band recurrent* neural coordination

| ROI Pair | Estimate | Std. Error | df | Lower CI | Upper CI | p-value |
| --- | --- | --- | --- | --- | --- | --- |
| Left anterior-Left anterior | 0.04 | 0.02 | 226.90 | -0.02 | 0.10 | 0.229 |
| Left anterior-Left temporal | 0.04 | 0.02 | 226.90 | -0.03 | 0.10 | 0.2499 |
| Left anterior-Medial anterior | 0.03 | 0.02 | 168.60 | -0.03 | 0.08 | 0.3853 |
| Left anterior-Medial central | 0.01 | 0.02 | 137.80 | -0.05 | 0.06 | 0.7078 |
| Left anterior-Right anterior | 0.08 | 0.02 | 226.90 | 0.02 | 0.14 | <0.001*** |
| Left anterior-Right temporal | -0.00 | 0.02 | 226.90 | -0.06 | 0.06 | 0.9622 |
| Left temporal-Left anterior | 0.01 | 0.02 | 226.90 | -0.05 | 0.07 | 0.7603 |
| Left temporal-Left temporal | 0.02 | 0.02 | 226.90 | -0.04 | 0.08 | 0.5404 |
| Left temporal-Medial anterior | -0.01 | 0.02 | 168.60 | -0.07 | 0.05 | 0.6867 |
| Left temporal-Medial central | 0.01 | 0.02 | 137.80 | -0.04 | 0.07 | 0.6867 |
| Left temporal-Right anterior | 0.01 | 0.02 | 226.90 | -0.05 | 0.07 | 0.7056 |
| Left temporal-Right temporal | 0.01 | 0.02 | 226.90 | -0.05 | 0.07 | 0.6867 |
| Medial anterior-Left anterior | 0.02 | 0.02 | 168.60 | -0.04 | 0.08 | 0.5404 |
| Medial anterior-Left temporal | -0.00 | 0.02 | 168.60 | -0.06 | 0.06 | 0.911 |
| Medial anterior-Medial anterior | 0.05 | 0.02 | 130.60 | -0.00 | 0.11 | 0.016* |
| Medial anterior-Medial central | 0.03 | 0.02 | 110.10 | -0.02 | 0.08 | 0.229 |
| Medial anterior-Right anterior | 0.06 | 0.02 | 168.60 | 0.00 | 0.12 | 0.0076** |
| Medial anterior-Right temporal | -0.01 | 0.02 | 168.60 | -0.07 | 0.05 | 0.6867 |
| Medial central-Left anterior | -0.00 | 0.02 | 137.80 | -0.06 | 0.05 | 0.9388 |
| Medial central-Left temporal | -0.01 | 0.02 | 137.80 | -0.06 | 0.05 | 0.8225 |
| Medial central-Medial anterior | 0.03 | 0.02 | 110.10 | -0.03 | 0.08 | 0.3433 |
| Medial central-Medial central | 0.01 | 0.02 | 95.00 | -0.04 | 0.07 | 0.5897 |
| Medial central-Right anterior | 0.02 | 0.02 | 137.80 | -0.03 | 0.08 | 0.4355 |
| Medial central-Right temporal | -0.02 | 0.02 | 137.80 | -0.07 | 0.04 | 0.5404 |
| Right anterior-Left anterior | 0.04 | 0.02 | 226.90 | -0.02 | 0.10 | 0.229 |
| Right anterior-Left temporal | 0.03 | 0.02 | 226.90 | -0.03 | 0.10 | 0.2914 |
| Right anterior-Medial anterior | 0.06 | 0.02 | 168.60 | 0.00 | 0.12 | 0.0061** |
| Right anterior-Medial central | 0.01 | 0.02 | 137.80 | -0.04 | 0.07 | 0.6867 |
| Right anterior-Right anterior | 0.09 | 0.02 | 226.90 | 0.03 | 0.15 | <0.001*** |
| Right anterior-Right temporal | -0.02 | 0.02 | 226.90 | -0.09 | 0.04 | 0.4834 |
| Right temporal-Left anterior | 0.01 | 0.02 | 226.90 | -0.06 | 0.07 | 0.798 |
| Right temporal-Left temporal | 0.03 | 0.02 | 226.90 | -0.04 | 0.09 | 0.3853 |
| Right temporal-Medial anterior | 0.02 | 0.02 | 168.60 | -0.04 | 0.08 | 0.4834 |
| Right temporal-Medial central | -0.01 | 0.02 | 137.80 | -0.07 | 0.04 | 0.6867 |
| Right temporal-Right anterior | 0.03 | 0.02 | 226.90 | -0.03 | 0.09 | 0.3433 |
| Right temporal-Right temporal | 0.00 | 0.02 | 226.90 | -0.06 | 0.07 | 0.911 |

Note. p-values are FDR-corrected.
