## Appendix D for "How do we align in good conversation? Investigating the link between interaction quality and multimodal interpersonal coordination"

### Appendix D: Does interpersonal coordination predict relational outcomes?

Communication Accommodation Theory frames interpersonal alignment as socially consequential (Giles et al., 1991), whereas proponents of the interactive-alignment model have suggested that neural coordination may provide a suitable indicator of interpersonal alignment across multiple levels of representation (Hasson et al., 2012; Menenti et al., 2012). This led us to ask, *a posteriori*, whether the inter-brain relationships that emerged as a meaningful marker of interaction quality would account for variance in the relational outcomes of different dyads’ interactions, as indexed by their mutual affiliation scores collected at the end of the experiment. We chose to focus on alpha-band, concurrent inter-brain relationship based on our previous results showing a robust association with PIQ, and on F0 and loudness turn-level synchrony given prior work indicating that these cues meaningfully index relational outcomes (De Looze et al., 2014). We applied PCA to the cross-brain, ROI-level GCMI values to reduce predictor dimensionality (from 36 ROI pairs). This step yielded a principal component that explained 29.3% of the variance in the data (with the second component accounting for only 9.1%). We used the first principal component as our main dual-brain predictor (PC_IB_), representing the major axis of systematic variation in concurrent inter-brain relationship. A multiple regression model of each dyad’s mutual affinity score as a function of their mean PC_IB_, PIQ, F0 synchrony, and loudness synchrony did not detect any significant effect, and a marginally significant effect of PIQ (PC_IB_: $F\left[ 1,19 \right]=2.46$, $p=.133$; PIQ: $F\left[ 1,19 \right]=3.89$, $p=.063$; $F0 sync: F\left[ 1,19 \right]=0.03$, $p=.856$; Loud. sync: $F\left[ 1,19 \right]=3.40$, $p=.081$). While non-significant, these exploratory statistics show some expected trends: dyads who had better conversations (i.e., higher PIQ) reported better mutual affinity, and dyads with greater mutual affinity synchronized their loudness better. Note, however, that this correlational analysis relies on only 24 datapoints (one mutual affinity score per dyad). Exploratory data analysis (Figure S2) suggests that, overall, mutual affinity scores increase with increasing cross-brain mutual information. It is therefore
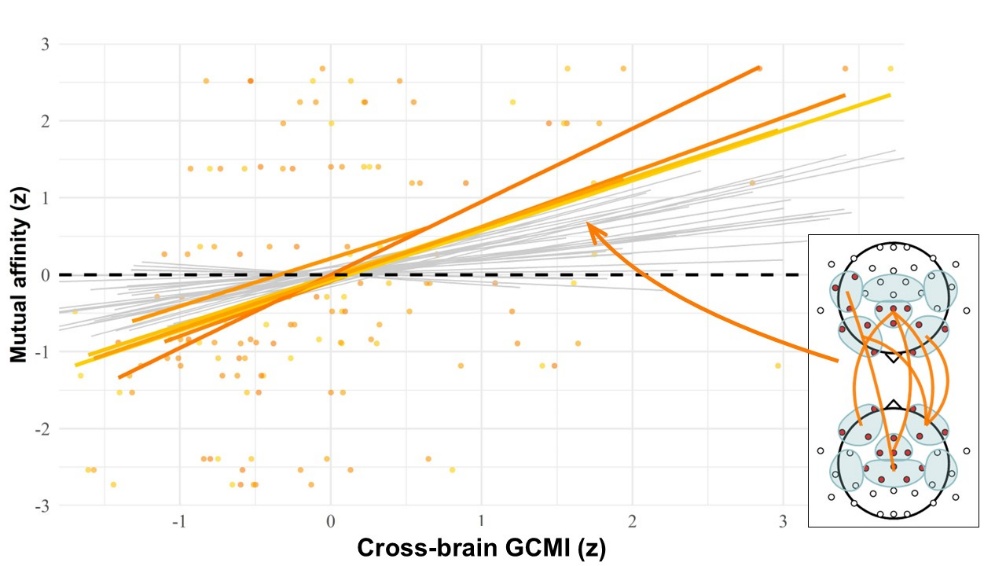
likely that our sample is underpowered for this exploratory analysis.

**Figure S2:** Exploratory association between neural coordination (alpha-band and concurrent) and relational outcomes (mutual affinity). Regression lines for all ROI pairs are shown in grey. Six ROI pairs with the strongest positive associations are highlighted in color to aid visualization; this subset is illustrative, and the choice to display six pairs, as well as the color coding, is arbitrary and does not reflect a statistical threshold. The overall pattern indicates a positive association between mutual affinity and inter-brain coupling, especially across fronto-medial regions.
